# Dual role of the receptor kinase FERONIA in regulating tissue mechanics and growth

**DOI:** 10.1101/2025.08.28.672783

**Authors:** Elise Muller, Marc Ropitaux, Gaëlle Durambur, Valentin Laplaud, Leon Gebhard, Arnaud Lehner, Stéphanie Drevensek, Arezki Boudaoud

## Abstract

During morphogenesis, growing organisms adapt to internal and external mechanical stress. In plants, the receptor kinase FERONIA is involved in a broad range of responses to perturbations of the cell wall, but it is unclear which cues are sensed and how this sensing controls growth. Here, we addressed these questions in Marchantia vegetative propagules. We combined culture in microfluidic chips to quantify growth and mechanical properties of propagules, perturbations with osmotic treatments, characterisation of cell wall polysaccharides, and a mathematical model of cell wall expansion that incorporates sensing. We found that FERONIA independently regulates tissue mechanical properties and growth rates. We propose that the regulation of growth by the FERONIA pathway relies on both a positive feedback from elastic deformation of the wall and a negative feedback from wall expansion. Altogether, we expect our quantitative framework to be broadly relevant to the investigation of how mechanical cues guide development.

## Introduction

The growth of a plant cell is driven by a high inner pressure, known as turgor pressure, and is limited by the surrounding cell wall, a stiff extracellular matrix mostly made of intertwined polysaccharides. Turgor-generated cell wall tension is relaxed by remodelling of the polysaccharide network or by integration of new polysaccharides (*1, 2*). Relaxation notably involves cell wall responses to the mechanical stress associated with tension. Responses may be “passive”, corresponding to elastic stretching of the polysaccharides or to plastic (irreversible) deformation arising from polysaccharides sliding along each other (*3*). However, the robustness and adaptability of plant growth in a fluctuating environment require “active” responses, which may be triggered by local mechanical stress or strain (*4*). Forces applied to cell walls are perceived by several classes of mechanosensors and/or sensors of cell wall integrity. Here, we focus on the role in growth of *FERONIA*, a salient member of the *Catharanthus roseus* receptor-like kinase 1-like (*Cr*RLK1L) family. *FERONIA* is involved in fertility, immunity, responses to biotic and abiotic stress, as well as in growth and development (*5*), which makes it difficult to study its specific function in growth.

Growth control by *FERONIA* was characterised in various organs and environmental conditions, leading to the conclusion that *FERONIA* is a context-dependent regulator (*6*). On the one hand, *FERONIA* may inhibit growth, as demonstrated by Arabidopsis *feronia* mutants (At*fer*), which display overgrowth of the pollen tube within the embryo sac (*7*) or bigger seeds (*8*). The secreted peptide Rapid Alkalinization Factor (RALF) activates AtFERONIA to suppresses cell elongation (*9*) or to inhibit vacuolar expansion (*10*). On the other hand, *FERONIA* may promote growth, as shown by the reduced size of At*fer* hypocotyls (*11, 12*), of rosettes (*13, 14*), or of root hairs and trichomes (*14*). Mechanosensing by *FERONIA* was also found to limit growth fluctuation in Arabidopsis roots (*15*). The role of *FERONIA* in growth regulation seems to be evolutionary conserved in land plant (*16*). A first study in the model liverwort *Marchantia polymorpha* reported a reduced thallus (vegetative body) in *feronia* mutants (Mp*fer*) (*16*).

Cell wall integrity sensors such as *FERONIA* mediate responses to cell wall damage (*17*). The nature of the activating signal and downstream signalling are still under investigation. Integrity sensors are mainly known to bind to wall-derived signalling molecules or to cell wall components, but may also act as a scaffold for cell wall proteins and peptides. In particular, FERONIA binds to RALF peptides (*5, 9*) and to pectins (preferentially de-methylesterified pectins) (*18, 19*). *FERONIA* interacts with co-receptors of the LRE-LIKE-GPI-AP family (LLG1 and LLG2) (*20*) and forms complexes with the extracellular LEUCINE-REACH REPEAT EXTENSINS (*21, 22*), that are also involved in cell wall integrity sensing. *FERONIA* responds to pathogen-associated molecular patterns, leading to cross-talks with immunity (*23*). It was also hypothesised that integrity sensors could act as true mechanosensors and respond to deformations of the membrane-cell wall continuum (*24*). This hypothesis is supported by reduced calcium signalling in At*fer* (*15*) upon mechanical bending and by the partial rescue of At*fer* by osmotic treatments to reduce mechanical stress (*25*). To assess what mechanical signals may activate *FERONIA* and how this contributes to growth regulation, we combine modelling and experiments. Indeed, the use of modelling has been instrumental to identify rules underlying the alignment of cortical microtubules in aerial organs (*26–28*) or strain-stiffening during seed growth (*29*).

After activation, FERONIA may trigger several downstream signalling pathways. It modifies calcium signalling, ROS production, apoplasmic pH, hormonal signalling, actin regulation, and vesicular trafficking, in addition to transcriptional regulation (*20, 30*). FERONIA regulates kinases as MARIS, a PTI-like protein involved in cell wall integrity of tip-growing cells – Arabidopsis pollen tube and Marchantia rhizoids (*31–33*).

Many phenotypes of At*fer* mutants point an alteration of the mechanical properties of cells and tissues (*22*). At*fer* mutants show reduced apoplasmic pH (*9*) and altered lignin composition (*34*). Cell bursting in tip-growing cells (*14, 16*) and in diffusely-growing pavement cells (*25*) indicates weaker cell wall or higher turgor in *fer* mutants, compared to wild-type. Indentation-based measurements show lower apparent modulus in At*fer* roots subject to salt stress and in Mp*fer* thalli, suggesting weaker cell wall or lower turgor than in wild-type. A mutant of another *Cr*RLK1L member, *THESEUS*, displays a decreased Brillouin elastic contrast (*35*). Here, we independently quantified turgor pressure and tissue elasticity to fully assess their regulation by *FERONIA*.

Altogether we aimed at building a quantitative framework to study the role of *FERONIA* in the regulation of tissue mechanics and growth, which led us to identify potential mechanical signals that may activate *FERONIA*. We took advantage of many features of *Marchantia polymorpha*. The *Cr*RLK1L has a single member Mp*FER*. Marchantia may reproduce vegetatively by producing a large number of genetically identical gemmae. Following immersion in water, gemmae germinate and start to grow (*36*) and can be cultivated in microfluidic chips (*37*). We thus investigated the role of *FERONIA* in gemmae growth and morphogenesis.

## Results

### *FERONIA* promotes and patterns growth in Marchantia gemmae

Previous work showed that Mp*fer* mutants have reduced cell and thallus sizes after several days or weeks of growth (*16*). To test whether this expands to early development, we recorded the first day of gemmae life in a microfluidic chip continuously supplied with liquid growth medium. The chip consists of a PDMS growth chamber, decorated with spacers – to isolate individual gemmae – and sealed with a glass slide (Fig. 1-A and B); we found that gemmae grow similarly in chips and outside of chips (*37*). Gemmae growth can be divided into two phases. A first rest phase shows almost no growth and lasts for a duration *T*_*start*_ of a few hours (one to ten hours), spanning imbibition (uptake of water upon immersion of gemmae) to germination (start of growth). The ensuing growth phase is characterised by an equilibrium growth rate *G*_*eq*_, with some fluctuations around this average (Fig. 1-C). Two independent Mp*fer* lines, *fer*-2 and *fer*-3 (*16*), exhibit both a late germination and a low equilibrium growth rate, compared to wild-type (Fig. 1-D to F). Thus, *FERONIA* promotes germination and growth, at gemma scale.

**Figure 1:**
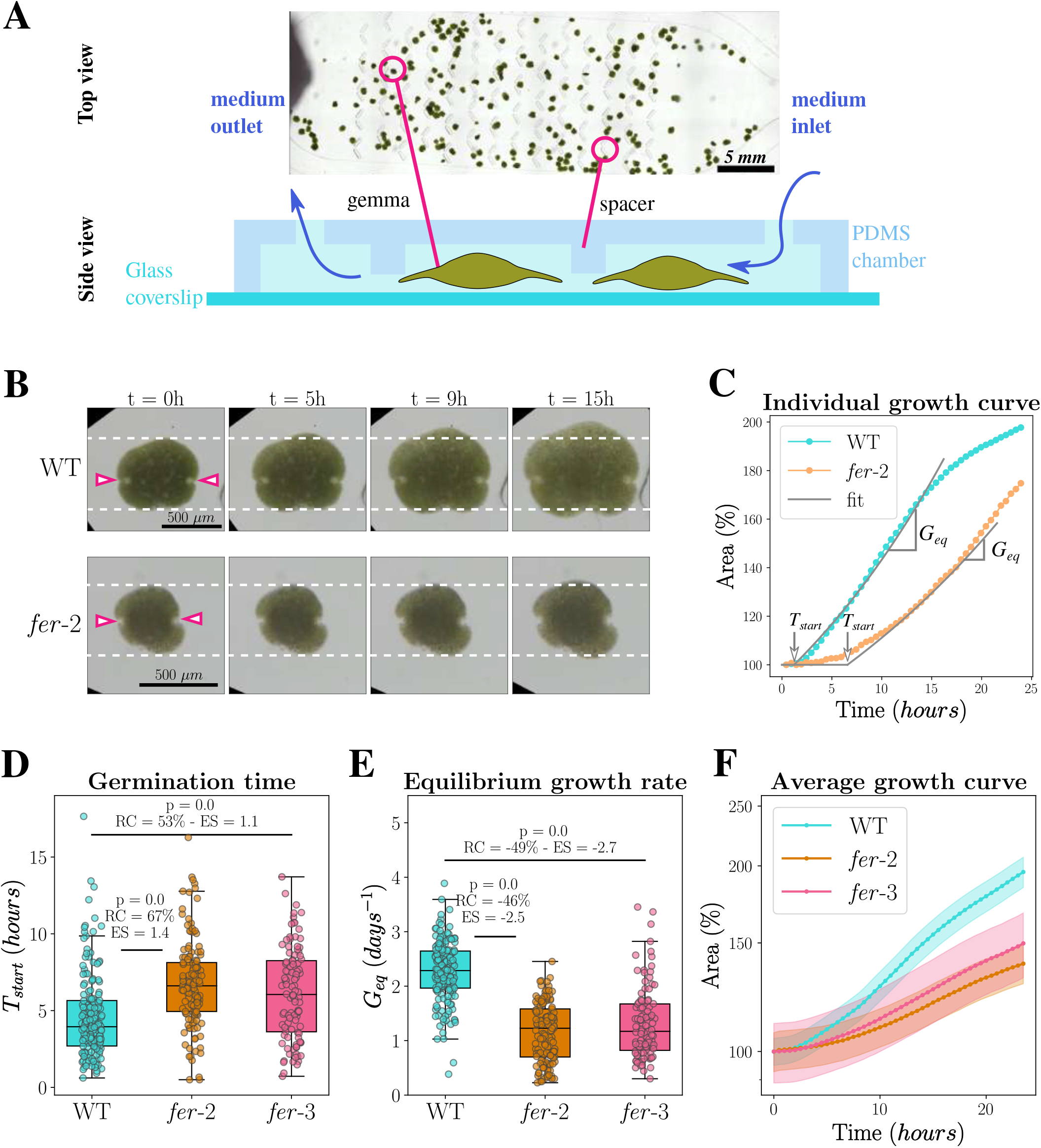
*FERONIA* promotes gemmae growth. (**A**) Microfluidic chip to grow gemmae. Upper panel: Top view of a microfluidic chip filled with gemmae. Medium inlet and outlet are on the sides. One gemma and one trap (spacer) are indicated with pink circles. The scale bar is 5 *mm* long. Lower panel: Sketch of the chip side view, one gemma and one spacer and indicated by pink lines. The (slow) flow of the culture medium is represented by blue arrows. (**B**) Bright field images of representative growing gemmae of WT and *fer*-2 at relevant time points (0 h, 5 h, 9 h and 15 h after imbibition, i.e. immersion in water). Meristematic regions (notches) are indicated by pink arrow heads. The dotted white lines are guides to observe gemmae growth. The scale bar is 500 µm long. (**C**) Representative individual growth curve of WT and *fer*-2 over the 24 first hours of growth after imbibition. The grey lines are the delayed exponential fit for each genotype. Characteristic parameters for growth are represented on the curves: the germination starting time *T*_*start*_ and the equilibrium growth rate *G*_*eq*_. (**D-F**) Growth of parameters of WT and mutants – *fer*-2 and *fer*-3 –. (**D**-**E**) Box plot and scatter plot of the germination starting time *T*_*start*_ (**D**) and equilibrium growth rate *G*_*eq*_ (**E**). The top and bottom of the box correspondto the lower and upper quartiles. (**F**) Average area (relative to initial area) as a function of time. The shaded areas correspond to the 95% confident interval. WT: n = 178 individuals and rep. = 5 replicates. *fer*-2: n = 141 and rep. = 3. *fer*-3: n = 104 and rep. = 3.

Next, we investigated whether *FERONIA* controls spatial differences in growth rate, notably because different functional domains make up a gemma. For instance, each gemma normally contains two meristematic regions known as apical notches (*38*) (highlighted by pink arrows in Fig. 1-B), from which derive more differentiated tissues during gemma development. To isolate the contribution of proliferation from that of elongation, the number of dividing cells was assessed by S-phase nuclei stained using EdU incorporation at different time points after imbibition. Proliferation is restricted to apical notches, consistent with classical descriptions (*38*) (pink arrows in Fig. 2-A). In wild-type (WT), proliferation can start as early as 2 h after imbibition, while first dividing nuclei in *fer*-2 are present only from 8 h to 10 h after imbibition. Moreover, even after proliferation begins in *fer*-2 gemmae, we still observe a higher number of dividing nuclei in WT than in *fer*-2 (Fig. 2-B). Accordingly, *FERONIA* promotes early timing and high rate of proliferation.

**Figure 2:**
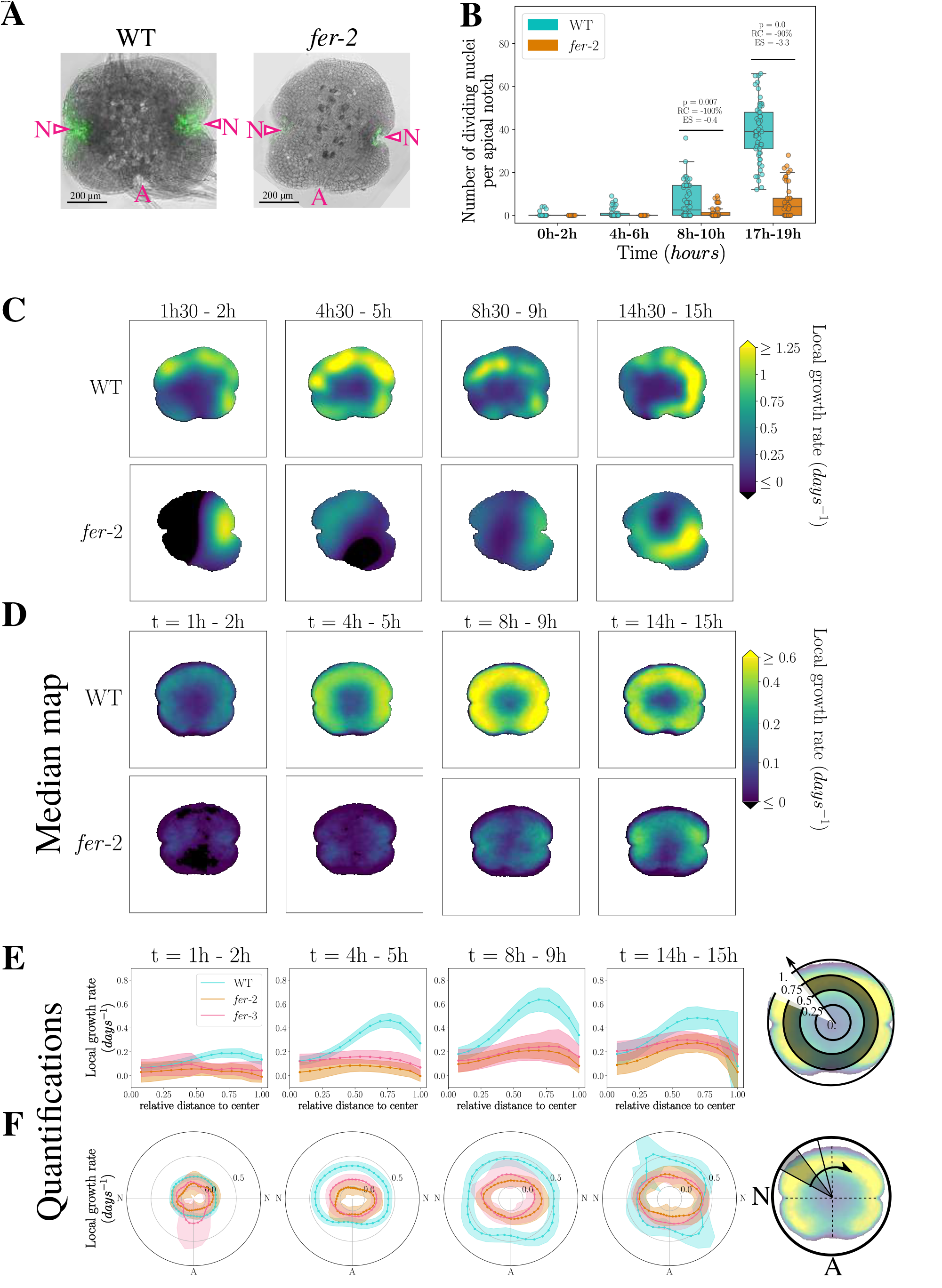
*FERONIA* regulates proliferation and local growth patterning. (**A-B**) Quantification of proliferation in WT and *fer*-2. (**A**) Representative merge confocal images of fluorescence intensity from EdU staining (green) and of bright field gemma view (gray) for WT and *fer*-2 for 17 h-19 h. Meristematic regions (notches) are indicated by pink arrow heads and letter N, while the former attachment point to the mother plant is indicated by the letter A. Scale bars are 200 µm in length. (**B**) Box plot and scatter plot of the number of dividing nuclei (stained with EdU) per notch for WT and *fer*-2 at 0 h-2 h, 4 h-6 h, 8 h-10 h and 17 h-19 h. WT: n(0 h-2 h) = 10, n(4 h-6 h) = 24, n(8 h-10 h) = 27, n(17 h-19 h) = 29, rep.= 2. *fer*-2: n(0 h-2 h) = 10, n(4 h-6 h) = 18, n(8 h-10 h) = 27, n(17 h-19 h) = 29, rep.= 2. (**C**) Local growth rate maps for representative individual gemmae of WT and *fer*-2. Local growth rate is quantified between two imaging time points (1 h30-2 h, 4 h30-5 h, 8 h30-9 h and 14 h30-15 h after imbibition) and is represented with a linear colour scale. (**D**) Median local growth rate maps for WT and *fer*-2. The median local growth rate is calculated over the individuals and over one hour at different time interval (1 h-2 h, 4 h-5 h, 8 h-9 h and 14 h-15 h after imbibition). It is represented with a symmetric logarithmic colour scale. WT: n = 61, rep. = 3. *fer*-2: n = 57, rep. = 3. (**E-F**) Radial and circumferential quantification of the local growth rate for WT, *fer*-2 and *fer*-3. (**E**) Mean local growth rate of the gemma surface at a given distance from the centre of the gemmae, averaged over all orientations (schematic on the right). (**F**) Mean local growth rate of the gemma surface in a given angular sector, averaged over all distances to the centre (schematic on the right). The shaded areas indicate the 95% confident interval. WT: n = 61, rep. = 3. *fer*-2: n = 57, rep. = 3.

We then considered the contribution of *FERONIA* to the regulation of elongation. Local, supra-cellular growth rate maps were generated by computing local displacement on growing gemmae (see (*39*)). Growth maps feature significant spatio-temporal variability (Fig. 2-C), although systematic differences between centre and periphery of gemmae can be suspected. To reveal such differences, we averaged the growth maps of about 60 individuals (see (*39*)) at different key time points (1-2 h which is before germination, 4 h-5 h at WT germination, 8 h-9 h at *fer*-2 germination, 14 h-15 h after germination) (Fig. 2-D). Starting from germination, a “growth ring” appears on WT gemmae (4 h-5 h time point), the intensity of which tends to increase (8 h-9 h time point) and finally reduces as the global growth rate diminishes (Fig. 2-D). To assess radial patterns of growth, we averaged local growth rate at a given distance from the centre of the gemma. Following germination, this radial average indeed shows a peak of growth rate at a relative distance of 75% from the centre (Fig. 2-E). For *fer*-2 and *fer*-3, the growth ring is almost absent (see Fig. 2-C and D, and radial quantification in Fig. 2-E). In the mutants, growth rate is maximal in the meristematic regions, appearing less affected there (Fig. 2-D). To assess circumferential patterns of growth, we averaged local growth rate at a given angle with respect to the line joining notches. This angular average is indeed higher at the angles corresponding to apical notches for *fer*-2 and *fer*-3 (Fig. 2-F). Angular averaging also shows that the local growth rate is diminished in *fer*-2 and *fer*-3 compared to WT, except at the angle of the apical notches, where mutants and WT local growth rates are the closest (Fig. 2-D and F). We note in particular that growth rate of *fer*-2 and *fer*-3 is reduced in regions without proliferation, meaning that *FERONIA* controls cell expansion. Altogether *FERONIA* regulates and patterns growth by independently promoting proliferation and elongation.

### *FERONIA* promotes and patterns gemmae turgor and stiffness, independently from growth regulation

In previous work, the lack of FERONIA activity was found to affect cell or tissue mechanics (*14,16, 18*), providing a potential explanation for growth defects in mutants. We thus wondered whether *fer* mutants gemmae exhibit mechanical defects. To address this question, we quantified the mechanical properties of WT and *fer*-2 gemmae, before germination (at 1 h after imbibition) and after (at 8 h after imbibition). We measured gemma elastic two-dimensional volumetric modulus (*E*_*gemma*_) and turgor pressure (*P*_*gemma*_) thanks to osmotic steps rapidly applied to the gemma in chip (Fig. 3-A and B). Equilibrium areas following step are extracted, and mechanical properties are deduced from the equilibration of water potential (detailed in (*39*)). As gemmae display the previously described strain-stiffening behaviour of plant walls – meaning that gemmae walls are stiffer as the deformation from the rest state increases – (*40*), different osmotic intensities were used to improve the determination of mechanical parameters (Fig. 3-A and (*39*)). Fig. 3-C and D show our results. First, *fer*-2 gemmae are softer than WT. Temporal variation of stiffness seems also affected by the lack of *FERONIA*, as a slight increase of the volumetric elastic modulus is observable in the WT between 1 h and 8 h, while the elastic modulus remains constant for *fer*-2 (Fig. 3-C). Similarly, *fer*-2 gemmae have a lower turgor pressure compared to WT, which tends to decrease after germination while it is maintained in WT (Fig. 3-D). So, at gemma scale, *FERONIA* promotes stiffness and a build-up of turgor.

**Figure 3:**
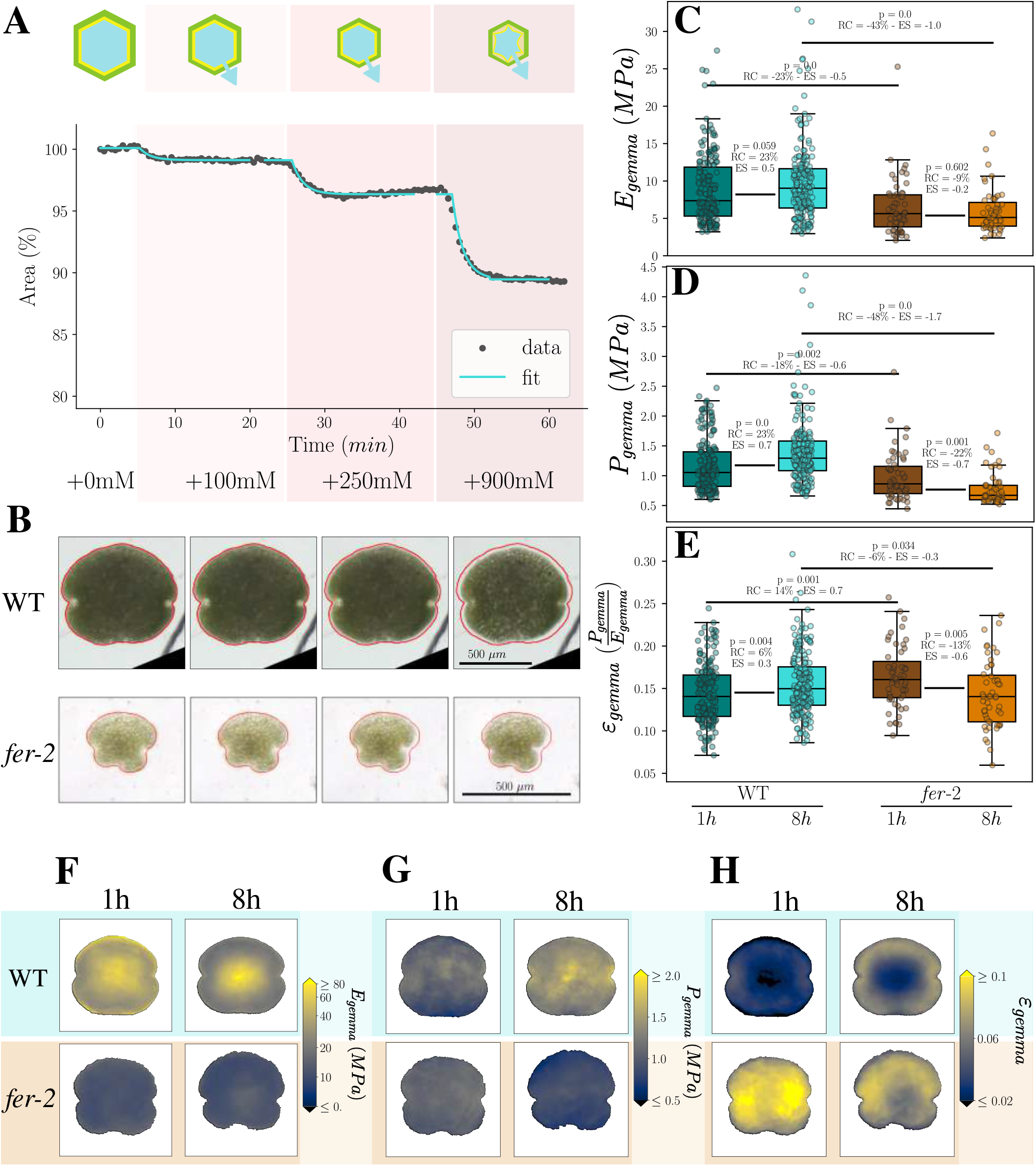
*FERONIA* regulates mechanical properties and their patterning. (**A**) Area of a gemma (% of initial area before osmotic steps) throughout an assay with several osmotic steps and fit of the predicted exponential decay to extract equilibrium values. Gemmae are subjected to three successive steps of +100 mM, +250 mM and +900 mM (the last leading to plasmolysis) of mannitol. The upper panel is a schematic of cell behaviours under these steps. The blue arrow represents net water flux. (**B**) Representative WT and *fer*-2 gemmae during osmotic steps (+0 mM, +100 mM, +250 mM and +900 mM). The red line is the initial contour of the gemma for each genotype and can be used as a visual guide for subsequent area reduction. Scale bars are 500 µm. (**C-E**) Box plot and scatter plot of measured mechanical properties at the gemma scale for WT and *fer*-2 gemmae at 1 h and 8 h after imbibition. (**C**) Volumetric elastic modulus *E*_*gemma*_, (**D**) turgor pressure *P*_*gemma*_ and (**E**) elastic deformation *ℰ*_*gemma*_ = *P*_*gemma*_/*E*_*gemma*_. WT: n(1 h) = 176, n(8 h) = 193, rep. = 4. *fer*-2: n(1 h) = 55, n(8 h) = 51, rep. = 3. (**F-H**) Maps of local mechanical parameters for WT and *fer*-2 gemmae at 1 h and 8 h after imbibition. Parameters are estimated locally from the median displacement during the osmotic steps. (**F**) Maps of local elastic modulus *E*_*gemma*_ with a symmetric logarithmic colour scale, (**G**) maps of turgor *P*_*gemma*_ with a linear colour scale and (**H**) maps of the elastic deformation, *ℰ*_*gemma*_ with a linear colour scale. WT: n(1 h) = 142, n(8 h) = 151, rep. = 3. *fer*-2: n(1 h) = 41, n(8 h) = 32, rep. = 3.

Next, we investigated whether such mechanical perturbation could be the cause of the growth deregulation in *fer* mutants. We tested whether gemmae at 8 h after imbibition grow in accordance with Lockhart’s law (*41*), according to which cell growth rate increases linearly with turgor pressure. Unexpectedly, gemma growth rate is not positively correlated to turgor (fig. S1-A). Our data rather show that growth rate negatively correlates with the volumetric elastic modulus *E*_*gemma*_, for both WT and *fer*-2 (fig. S1-B). We previously observed the same negative correlation at 30 h after imbibition, which leads to the hypothesis that, for gemmae, the elastic deformation *ℰ*_*gemma*_ = *P*_*gemma*_/*E*_*gemma*_ is the relevant mechanical factor to analyse growth (*37*). In our 8 h gemma data set, global growth rate is indeed positively correlated with elastic deformation *ℰ*_*gemma*_, this correlation being stronger than with *P*_*gemma*_ or *E*_*gemma*_ individually (Fig. 5-A). We thus considered the following equation to describe gemma growth (equation 1),

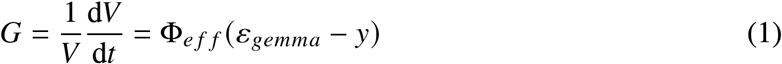

which defines the instantaneous growth rate *G* as the relative rate of volume *V* change, with Φ_*eff*_ the effective extensibility, *ℰ*_*gemma*_ the elastic deformation and *y* the deformation threshold for growth to occur (*G* is set to 0 if *ℰ*_*gemma*_ *< y*).

We tried to interpret our mechanical data in terms of growth using Equation 1. First, effective extensibility is predicted to be enhanced in *fer*-2 because it is softer than WT (Fig. 3-C), which should lead to faster growth. Second, *ℰ*_*gemma*_ is rather similar between WT and *fer*-2 at 8 h, and is even higher for *fer*-2 at 1 h, which also leads to faster growth. Both effects are in contrast with observations of slower growth in *fer*-2 than in WT (Fig. 1-C). Therefore, mechanical parameters cannot explain differences in growth between *fer*-2 and WT.

Because those measurements are global and might hide local mechanical features relevant for growth, we turned to the measurement of mechanical parameters at a smaller (though supracellular) scale. We analysed local deformations during the osmotic steps and interpreted them in terms of mechanical parameters to create average map of stiffness and turgor (Fig. 3-F and G, and detailed in (*39*)). While turgor maps seem rather homogeneous (Fig. 3-G), stiffness maps display a strong radial patterning, with the outer region being softer than the inner region (Fig. 3-F). This radial patterning is much stronger in WT than in *fer*-2 (Fig. 3-F). The local map of elastic deformation *ℰ*_*gemma*_ is also patterned in WT (especially at 8 h), which is reminiscent of local growth rate patterns at the same time (Fig. 3-H and Fig. 2-D). Patterning of local elastic deformation is almost absent in *fer*-2 (Fig. 3-H).

Altogether, *FERONIA* regulates both stiffness and turgor of gemmae, which might be locally associated to growth pattern but cannot explain global growth differences between *fer*-2 and WT.

### Indirect evidence that *FERONIA* and *MARIS* modulate cell wall extensibility

Given that growth regulation by *FERONIA* does not arise from mechanical regulation, we next hypothesised that *FERONIA* regulates extensibility. To test this hypothesis, we took advantage of our extensive dataset and analysed the relation between instantaneous growth rate averaged between 6 h and 8 h after imbibition and elastic deformation *ℰ*_*gemma*_ at 8 h. Following Equation (1), we fitted this relation to line, the slope of which corresponds to extensibility. We found that extensibility is higher in WT than in *fer*-2, although the spread of the data does not allow a strong conclusion (Fig. 5-A).

Mutation of *FERONIA* causes a relatively strong phenotype, affecting in particular the formation of the organs in which gemmae develop, the gemma cups (*16*), raising the possibility of maternal effects on gemmae. We therefore also considered MARIS, a kinase downstream of FERONIA, as under our growth condition gemma cups of *mri*-1 knock-down mutant are similar to those of the WT (*32*). We investigated whether growth and extensibility are affected in *mri*-1 mutant. *mri*-1 mutant displays reduced growth, similar to but weaker than in *fer*-2, with later germination and reduced equilibrium growth rate (fig. S2-A to C). Note that we use two controls (WT and WT-2) because the *mri*-1 mutant line was isolated by screening a cross between two accessions (*33*). Moreover, *mri*-1 local growth rate maps exhibit features similar to *fer*-2 and *fer*-3: a reduced growth ring compared to the WTs and the growth rate of meristematic regions is the least affected (fig. S2-D to F). We measured the mechanical parameters of *mri*-1 with osmotic steps and found minimal differences with wild-type (fig. S2-G to I). We computed local mechanical patterns in *mri*-1 and found them to be close to WT patterns (fig. S2-J to L). These observations uncouple mechanical patterning and growth patterning. Altogether, growth alterations in *mri*-1 cannot be ascribed to alterations of mechanical parameters, but potentially to altered extensibility. This defect in extensibility might be shared with *fer* mutants.

We thus looked for molecular effectors of extensibility that would be regulated by both *FERONIA* and *MARIS*. We took advantage of the recently published transcriptomes of *fer*-1 and *mri*-1 (*42*) and we sought whether there were cell-wall related genes among differentially expressed genes in both mutants. Transcripts related to xyloglucan:xyloglucosyl transferase activity are significantly enriched among the downregulated transcript (the corresponding GO:0016762 is significantly enriched with statistical significance *p* ≤ 0.001 in *fer*-1 and *p* ≤ 0.05 in *mri*-1). Enzymes of the xyloglucan endotransglucosylase/hydrolase (XTH) family are involved in the remodelling of xyloglucan, a core polysaccharides composing the plant cell wall. Thus, their down regulation suggests a reduced extensibility of mutant plants. Indeed, overexpression of XTH leads to stiffer hypocotyls (*43*), possibly explaining the lower stiffness in *fer*-2. Moreover, in Arabidopsis, mutants affected in xyloglucan remodelling enzymes (XHT family) often display growth phenotypes, which could be due to reduced extensibility (*44*) and mutants with virtually no xyloglucan have reduced measured extensibility (*45*).

To test whether xyloglucan is affected in *fer*-2, we analysed polysaccharides repartition in the cell wall of gemmae using immunolabelling (list of the monoclonal antibodies and their recognised epitopes are listed in table S2). We detected a decrease in the fluorescence associated to both the xyloglucan backbone (~−35% LM15, Fig. 4-A to C) and of galactosylated side-chains (~−24%, LM24 Fig. 4-D to F; ~−30% LM25, Fig. 4-G to I) in *fer*-2 mutant compared to WT. As Marchantia do not possesses xyloglucan fucosylated side chain, we did not used the CCRCM1 antibody recognising those patterns (*46*). Moreover, signal reduction seems to be specific to xyloglucan, as the other hemicelluloses tested, heteromannan with LM21, have similar labelling pattern and level of fluorescence between WT and *fer*-2 (Fig. 4-J to L). This results suggest changes either in xyloglucan content or in xyloglucan accessibility in the cell wall, which might explain part of the reduction of growth in *fer*-2.

**Figure 4:**
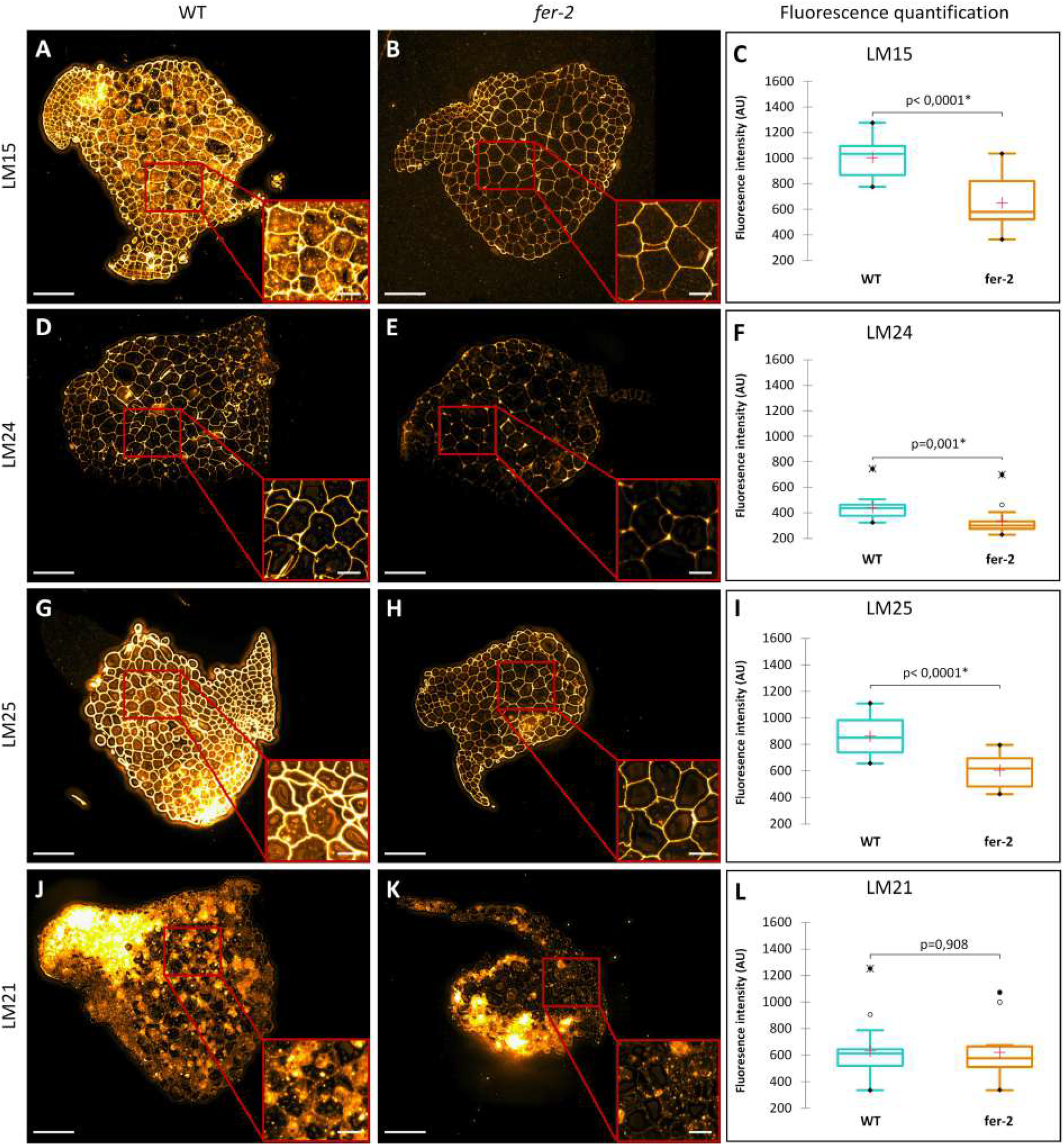
*FERONIA* affects cell wall hemicellulosic polymers distribution. (**A**,**D**,**G**,**J**) Immuno-cytochemical labelling of WT and (**B**,**E**,**H**,**K**) *fer*-2 mutant line gemmae. All samples were fixed, dehydrated, embedded and cut (2 µm thickness) before immunolabelling with primary antibodies: LM15 (Xylosylated xyloglucan (XXXG)), LM24 (Galactosylated xyloglucan (XXLG)), LM25 (Galactosylated xyloglucan (XXLG, XLLG)), and LM21 (β-(1→4)-mannan backbone epitope from heteromannan). (**C**,**F**,**I**,**L**) Box plots illustrating the fluorescence intensity. Each histogram represents the means of 14 biological replicates. Mann-Whitney tests were used; the start (*) represents de significant difference at level alpha = 0.05. The red crosses correspond to the means. The black circles correspond to extreme points and crossed-out black circles to extreme values excluded from the analysis. Scale bars are 100 µm for main images and 25 µm for magnified images.

**Figure 5:**
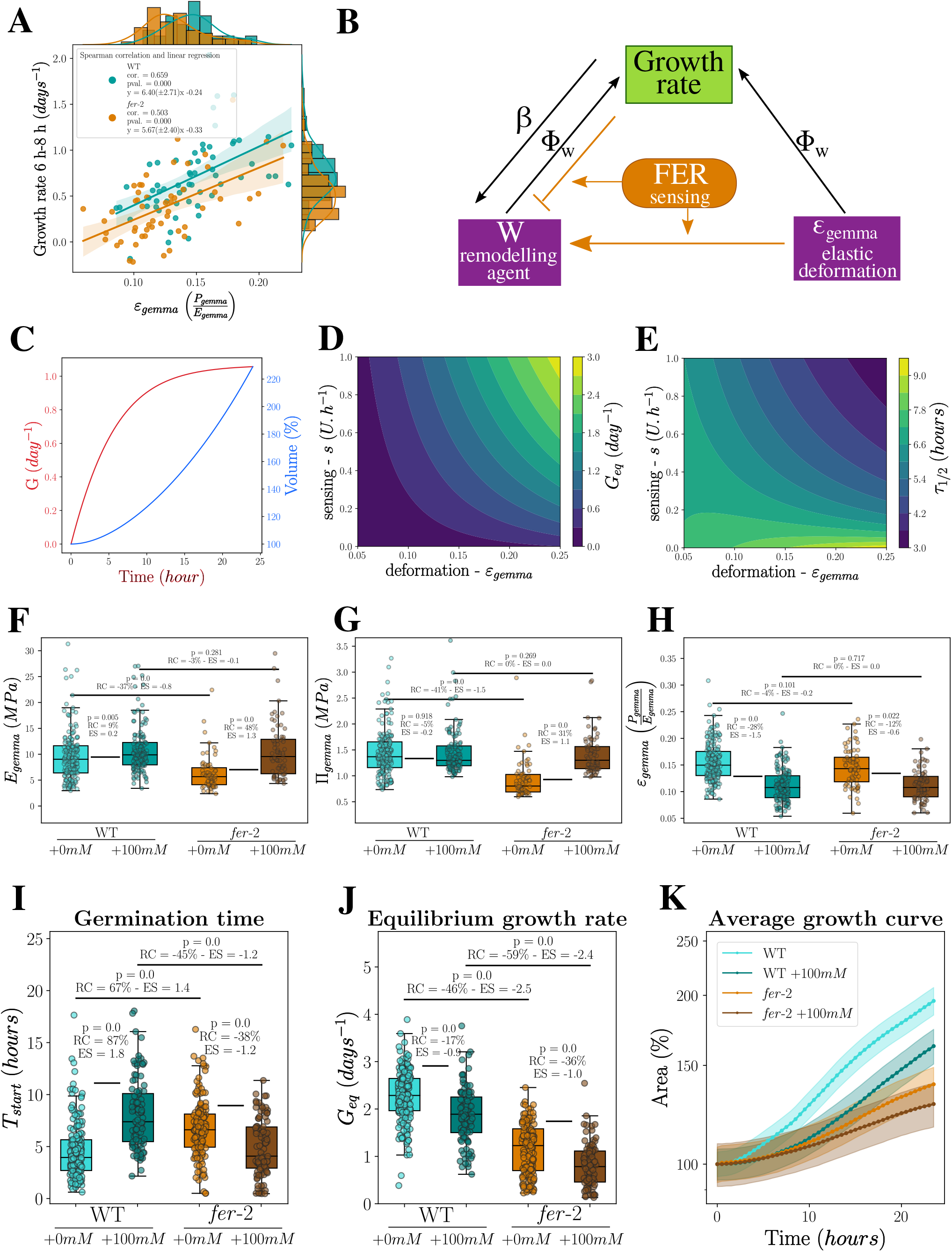
*FERONIA* responds to deformation and regulates extensibility. (**A**) Correlation analysis of the instantaneous growth rate averaged between 6 h and 8 h post imbibition with the elastic deformation *ℰ*_*gemma*_ for WT and *fer*-2. Scatter plot and regression line with 95% confident interval are represented as well as histograms per genotype for each variable. Correlations are quantified by the Spearman correlation coefficient (*cor*.) and a linear model is fitted to the data, which parameters are given with the 95% confident interval. WT: n = 62, rep. = 3. *fer*-2: n = 56, rep. = 3. (**B**) Schematic representation of the minimal mathematical model, describing the relationship between the growth rate, the elastic deformation (*ℰ*_*gemma*_), the unknown remodelling agent *W* and *FERONIA* sensing, and dependence on the autocatalytic rate *β* and the molar extensibility Φ_*w*_. (**C**) Instantaneous growth rate *G* and volume *V* over time as calculated by the model for 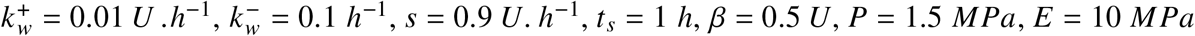 and *Y* = 0.01. (**D-E**) Phase diagram of the calculated growth parameters according to the toy model depending on sensing intensity *s* and on deformation *ℰ*_*gemma*_, for 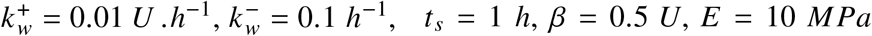 and *Y* = 0.01. (**D**) Calculated equilibrium growth rate *G*_*eq*_ and (**E**) and germination time *τ*_1/2_. (**F-H**) Box plot and scatter plot of the mechanical properties for 8 h after imbibition WT and *fer*-2. Plants were grown in reference medium with +0 mM or with +100 mM mannitol. (**F**) Osmotic pressure Π_*gemmae*_, (**G**) volumetric elastic modulus *E*_*gemma*_ and (**H**) elastic deformation *ℰ*_*gemma*_ are plotted for each condition. WT(+0 mM): n = 193, rep. = 4. WT(+100 mM): n = 136, rep. = 3. *fer*-2(+0 mM): n = 69, rep. = 6. *fer*-2(+100 mM): n = 70, rep. = 3. (**I-K**) Parametrisation of growth of WT as well as of *fer*-2 for plants grown in reference medium with +0 mM and with +100 mM mannitol. (**I**) Box plot and scatter plot of the germination starting time *T*_*start*_. (**J**) Box plot and scatter plot of the equilibrium growth rate *G*_*eq*_. (**K**) Average area over time (relative to initial area), the shaded areas correspond to the 95% confident interval. WT: n = 178, rep. = 5, WT(+100 mM): n = 99, rep. = 3, *fer*-2: n = 141, rep. = 3, *fer*-2(+100 mM): n = 94, rep. = 3.

Altogether, these data suggest that the FERONIA/MARIS (FER/MRI) pathway affects extensibility through xyloglucan levels and xyloglucan remodelling.

### A mathematical model of the regulation of extensibility by the FER/MRI pathway recapitulates growth phenotypes

In order to understand how *FERONIA* and *MARIS* may control growth through the regulation of extensibility, we built a minimal mathematical model based on the following considerations.

1. Growth rate *G* is classically described by the Lockhart equation, modified with the contribution of the elastic modulus *E*_*gemma*_ as in Equation (1) (*37, 41*).
2. We hypothesised that the effective extensibility is set by the activity of a putative remodelling agent that represents all potential remodelling agents, of surface concentration *W*. Thus, Φ_*eff*_ = Φ_*w*_*W*, where Φ_*w*_ is the agent molar extensibility; this decomposition was previously used to model pectin remodelling during pollen tube expansion (*47*). In the light of the indirect evidence that xyloglucan remodelling explains lower extensibility of *fer*-2 and *mri*-1 mutants, variations in agent concentration *W* might be ascribed to variations of the concentrations of enzymes involved in xyloglucan remodelling.
3. We assume that the remodelling agent is synthesised and integrated into the cell wall at a basal rate of 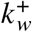 and a growth-dependent rate *β*, as proposed in the context protein synthesis during animal cell growth (*48*). The remodelling agent is degraded at a rate 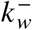 and diluted by wall expansion at rate *G*. Our core hypothesis is that *W* is affected by the activity of FERONIA and MARIS and by signalling through FERONIA/MARIS pathway, which is described by a function 𝒮 to be defined hereafter. The dynamics of agent concentration thus follows

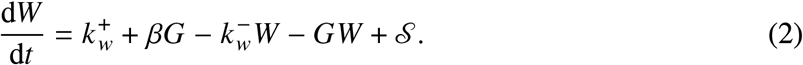
4. Quantification of growth shows that sensors affect both dynamics of growth (by speeding up germination) and equilibrium growth rate (by increasing it). We thus hypothesised that the effect of FERONIA/MARIS on extensibility is regulated by both a dynamical parameter, responsible for germination speed, and a static parameter, responsible for equilibrium growth rate. The minimal natural parameters are elastic strain (deformation *ℰ*_*gemma*_ = *P*_*gemma*_/*E*_*gemma*_) and irreversible strain rate (growth rate *G*) (*49*). We thus write the sensing function 𝒮 in the form

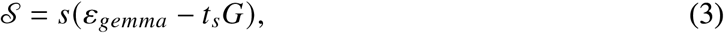

where *s* is the strength of the feedback and *t*_*s*_ a characteristic time scale to sense elongation. The signs of the two terms are constrained by experimental observations. In *fer*-2 compared to WT, sensing is reduced (*s* is smaller), deformation *ℰ* is roughly equivalent, and extensibility is reduced (growth is slower), hence a positive sign for the first term. In *fer*-2 compared to WT, the increase in extensibility is slower (germination is retarded), hence a negative sign for the second term.

We used this model (Equations 1-3) to predict germination time and equilibrium growth rate, and we compared these predictions to experimental observations (Fig. 1-D to F and fig. S1-C). We first determined plausible ranges for the parameters based on literature values (table S1) and checked predictions of *G*_*eq*_ and *T*_*start*_ are comparable in magnitude to their experimental values (model parameters exploration in fig. S1-D and E).

We then assessed consistency of model predictions with observations. Gemma volume is predicted to change slowly initially and then increases exponentially (Fig. 5-C), which resembles experimental behaviour (Fig. 1-C and F). Growth rate increases from 0 at the time corresponding to imbibition and reaches a plateau (Fig. 5-C), which is also observed in experiments (fig. S1-C), albeit with oscillations at later times. We analysed the model and solved it numerically (Supplementary Text). We confirmed that a decrease of sensing strength *s* reduces growth (decreases equilibrium growth rate *G*_*eq*_, Fig. 5-D) and retards germination (decreases germination time *τ*_1/2_, Fig. 5-E), as observed when mutating *FERONIA* (Fig. 1-D and E) or *MARIS* (Fig. S2-A and B).

Altogether, sensing both elastic deformation and growth by the FER/MRI pathway is sufficient to explain gemmae growth phenotypes.

### The model predicts responses of the FER/MRI pathway to osmotic pressure

We next considered mechanical perturbations to test our model. We varied the value of the elastic deformation *ℰ*_*gemma*_ in the model and we examined changes in equilibrium growth rate and germination time (see below). Experimentally, we sought to affect elastic deformation using long-term, mild osmotic treatments by addition of 100 mM mannitol in the culture medium. Because gemmae might osmoregulate, i.e. increase their content in osmolytes to compensate for the external increase so as to maintain their turgor pressure, we first quantified osmotic pressure under permanent osmotic treatment. Osmotic pressure is increased in treated *fer*-2 (compared to untreated), indicating osmoregulation, while the WT shows no osmoregulation (Fig. 5-G). To fully assess the mechanical state of gemmae, we also measure the elastic modulus. *fer*-2 appears stiffer under treatment, while WT shows a much reduced increase in stiffness (Fig. 5-F). We note that mechanical properties of WT gemmae are less affected by the treatment than *fer*-2 gemmae, consistent with the notion that FER is involved in mechanical responses. Finally we examined elastic deformation *ℰ*_*gemma*_ in these experiments. *ℰ*_*gemma*_ is diminished in similar proportions under treatment in WT and *fer*-2 (Fig. 5-H).

We then compared model predictions for germination time and growth rate with observations.

The equilibrium growth rate *G*_*eq*_ is predicted to decrease when elastic deformation is reduced, regardless of the value of the sensing parameter *s*, meaning that this behaviour is expected for all genotypes (equation S20 and Fig. 5-D). As predicted, if gemmae are continuously treated with +100 mM (Fig. 5-K for experimental data), the equilibrium growth rate *G*_*eq*_ decreases in WT as well as in *fer*-2 and *mri*-1 (Fig. 5-J and fig. S1-G and H). In contrast, changes in germination time (quantified by *τ*_1/2_ in the model) according to elastic deformation are predicted to depend on the sensing parameter *s*. If *s* is high (typically 0.9 *U. h*^−1^, to mimic WT), germination is predicted to be delayed when elastic deformation is reduced, while if *s* is low (typically 0.05 *U. h*^−1^, to mimic *fer*-2), germination is predicted to be advanced when elastic deformation is decreased (equation S21 and Fig. 5-E). In agreement with predictions, germination is retarded under treatment in WT while it is accelerated in *fer*-2 (Fig. 5-I). Interestingly, *mri*-1, which has an overall milder phenotype than *fer*-2, is expected to have an intermediate level of sensing and so an intermediate behaviour for germination time. As expected, germination time is not affected by the treatment in *mri*-1 (fig. S1-F). Altogether, those results support the hypothesis that extensibility is positively regulated by elastic deformation and negatively regulated by growth rate through the FER/MRI pathway.

### The FER/MRI pathway regulates variability of growth

It was proposed that responses to mechanical signals provide feedback loops to ensure developmental robustness (*50–52*). Interestingly, our mathematical model of the FER/MRI pathway involves several feedback loops, suggesting a function in the regulation of growth variability. Indeed, mutation of *FERONIA* in Arabidopsis increases local growth variability in the root (*15*). We therefore investigated the regulation of growth variability of gemmae by *FERONIA*.

We first reexamined our mathematical model regarding variability. We estimated the variability of the germination time *τ*_1/2_ as its relative sensitivity to the initial value of the remodelling agent *W*; we defined the variability of the equilibrium growth rate *G*_*eq*_ as its relative standard deviation under noise in the production of the remodelling agent (supplementary model). The model predicts an increase of variability in germination characteristic time when sensing is decreased and, to a lesser extent, when elastic deformation *ℰ*_*gemma*_ is decreased (Fig. 6-A). The same trends are predicted for the variability in equilibrium growth rate *G*_*eq*_ (Fig. 6-B). These predictions are consistent with the idea that *FERONIA* promotes growth robustness through feedback loops.

**Figure 6:**
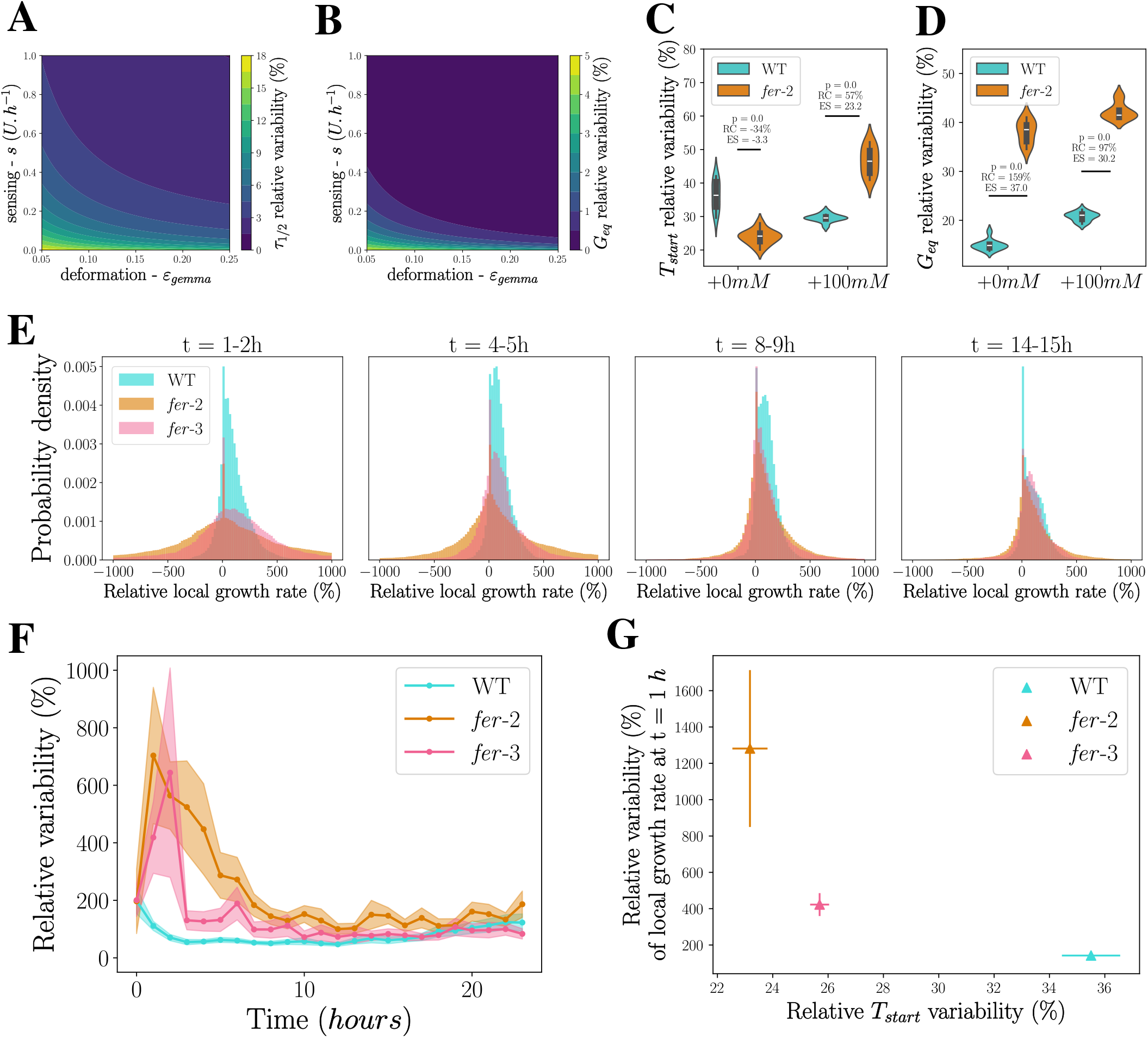
*FERONIA* regulates growth variability. (**A-B**) Relative variability of the growth parameters of the mathematical model according to the sensing parameter *s* and the elastic deformation *ℰ*_*gemma*_ for 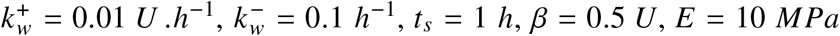 and *Y* = 0.01, of the germination time *τ*_1/2_ (given by 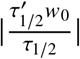) (**A**) and of the equilibrium growth rate *G* (given by 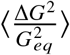) (**B**). (**C-D**) Violin plot of the relative variability in experiments (estimated by hte average absolute deviation normalised by the mean) of *fer*-2 and WT at +0 mM and +100 mM for *T*_*start*_ (**C**) and *G*_*eq*_ (**D**). WT: n = 178, rep. = 5, WT(+100 mM): n = 99, rep. = 3, *fer*-2: n = 141, rep. = 3, *fer*-2(+100 mM): n = 94, rep. = 3. (**E**) Probability density of local growth rate (normalised by its median) for WT, *fer*-2 and *fer*-3 at different times after imbibition. (**F**) Quantification of the variability (AAD over the median) of the local growth rate during 24h of growth, for WT, *fer*-2 and *fer*-3, the shaded areas represent the 95% confident interval. WT: n = 61, rep. = 3. *fer*-2: n = 57, rep. = 3, *fer*-3: n = 50, rep. = 3. (**G**) Relative variability of the local growth rate at 1 h (mean AAD over the median) as a function of the relative variability of *T*_*start*_ for WT, *fer*-2 and *fer*-3. Error bars are 90% confidence interval. WT: n = 61, rep. = 3. *fer*-2: n = 57, rep. = 3, *fer*-3: n = 50, rep. = 3.

We then tested these predictions and quantified experimental variability of the germination time *T*_*start*_ and equilibrium growth *G*_*eq*_ for wild-type plants, *fer* mutants and *mri*-1 under +0 mM and +100 mM osmotic conditions (Fig. 6-C and D and fig. S4-A to D). Considering the equilibrium growth rate *G*_*eq*_, it is clear that a decrease of FER/MRI pathway activity induces a higher variability. Indeed, both *fer*-2 and *fer*-3 have a higher variability than WT (Fig. 6-D and fig. S4-B). The same results hold for *mri*-1 (fig. S4-D). The variability of the equilibrium growth rate is also increased by a reduced elastic deformation (meaning under +100 mM osmotic treatment) in all genotypes — WT, *fer*-2 and *mri*-1 (Fig. 6-D and fig. S4-D). All observations are in line with predictions, confirming the role of the FER/MRI pathway, together with elastic deformation, in reducing the variability of the equilibrium growth rate *G*_*eq*_. Variability of the germination time *T*_*start*_ is also increased in the *fer*-2 mutant compared to WT, but only for gemmae growing under +100 mM osmotic stress (Fig. 6-C). For the reference culture conditions (+0 mM), the variability of the germination time *T*_*start*_ seems to decrease in the *fer*-2 mutant (Fig. 6-C and fig. S4-A). These trends are confirmed by the analysis of *mri*-1: with +100 mM osmotic stress *mri*-1 gemma germination is more variable than WTs, while *mri*-1 germination time variability is in the range of WTs without osmotic stress (fig. S4-C). Experimental results disagree with predictions in normal culture medium and agree under osmotic treatment. To sum up, *FERONIA* promotes a decrease of variability of gemmae growth, with the notable exception of germination time under normal turgor pressure.

To understand why the variability of germination time is lower in *fer* mutants than in WT in the reference culture medium (+0 mM osmolytes), we considered spatial variations of the growth rate. We noticed a broader distribution of local growth rate (normalised by the mean) in *fer*-2 and *fer*-3 mutants, compared to WT (Fig. 6-E). When plotting the relative width of growth distributions as a function of time (Fig. 6-F), it appeared that there is an initial peak of variability before germination in the two *fer* mutants. Two important remarks follow. First, at local scale, even in the reference medium (+0 mM osmolytes), mutating *FERONIA* increases variability. Second, there is a correlation between high local growth variability before germination and low global germination time *T*_*start*_ variability. This correlation is visible for the different mutants (Fig. 6-G). One interpretation of this correlation is that, in the absence of FERONIA, there is a higher stochasticity of local growth rate which averages on the whole gemmae and leads to a more reproducible behaviour, while, if FERONIA is present, this local stochasticity is diminished and the whole gemmae behaviour is more coordinated. So, in WT, there is less averaging and the global behaviour appears less robust. Indeed, spatiotemporal averaging reduces organ scale variability, as shown by models developed for Arabidopsis sepals (*53*).

## Discussion

We provided evidence that FERONIA and the downstream kinase MARIS regulate growth of Marchantia gemmae, in line with previous studies highlighting their role in growth of Arabidopsis and Marchantia (*7–11, 14, 16, 31, 32*). We showed that the FER/MRI pathway positively regulates germination of gemmae. *FERONIA* was also involved in germination of Arabidopsis seeds, although it was found to promote germination arrest in response to abscissic acid (*54*). We found that *FERONIA* promotes both cell proliferation and cell expansion and that *FERONIA* is necessary for the patterning of growth in gemmae. In At*fer* roots, proliferation defects can be observed (*55*) and defective growth patterns have been quantified (*15*).

We demonstrated a significant effect of *FERONIA* on the mechanical properties of gemmae. *FERONIA* is also required to maintain a radial pattern of elastic modulus, with softer gemma periphery. This is consistent with indentation-based measurements in *feronia* mutants showing either reduced stiffness or reduced turgor, in Arabidopsis (*18*) and in Marchantia (*16*), and with Brillouin microscopy-based measurements in At *theseus* mutant, another *Cr*RLK1 member (*35*). Here, we independently measured elastic modulus and turgor pressure, and we showed that *FERONIA* promotes build-up of both stiffness and turgor. In Arabidopsis roots, *FERONIA* controls the expansion of the vacuole (*10*), and so is potentially involved in turgor regulation.

We found that, in a given genotype, growth rate correlates with the elastic deformation *ℰ*_*gemma*_, as proposed in a chemorheological modelling framework for wall expansion (*47*) and partially tested in Marchantia (*37*). Nevertheless, elastic deformation is unaffected (or even increased) in *fer*-2 gemmae, which does not explain their slower growth compared to wild-type. Inspired by the proposal that strain-stiffening patterns growth domains (*56*), it is tempting to associate radial patterns of growth and of elastic modulus as both patterns are lost in *fer*-2. However, elastic modulus and growth are uncoupled in *mri*-1, in which only the growth pattern is affected compared to wild-type.

We thus uncovered a dual role for *FERONIA*: it independently controls mechanical properties (stiffness, turgor, elastic deformation) and growth. This implies that *FERONIA* regulates extensibility, i.e. the ability of the cell wall to expand under a given tension. In *mri*-1, growth is affected like in *fer*-2, whereas most mechanical defects observed in *fer*-2 are absent. This suggests that the regulation of extensibility is *MARIS*-dependent while the regulation of mechanical properties depends on other pathways downstream of *FERONIA* (Fig. 7-A). In the context of pollen tube growth regulation, it was shown that *MARIS* belongs to only part of the regulation pathways downstream of the pollen tube major *Cr*RLK1 *ANXUR1/2* (*31, 57, 58*). *FERONIA* was associated with multiple downstream effectors — pH, calcium, or ROS (*5*) — that might be responsible for direct cell wall remodelling and so extensibility increase.

**Figure 7:**
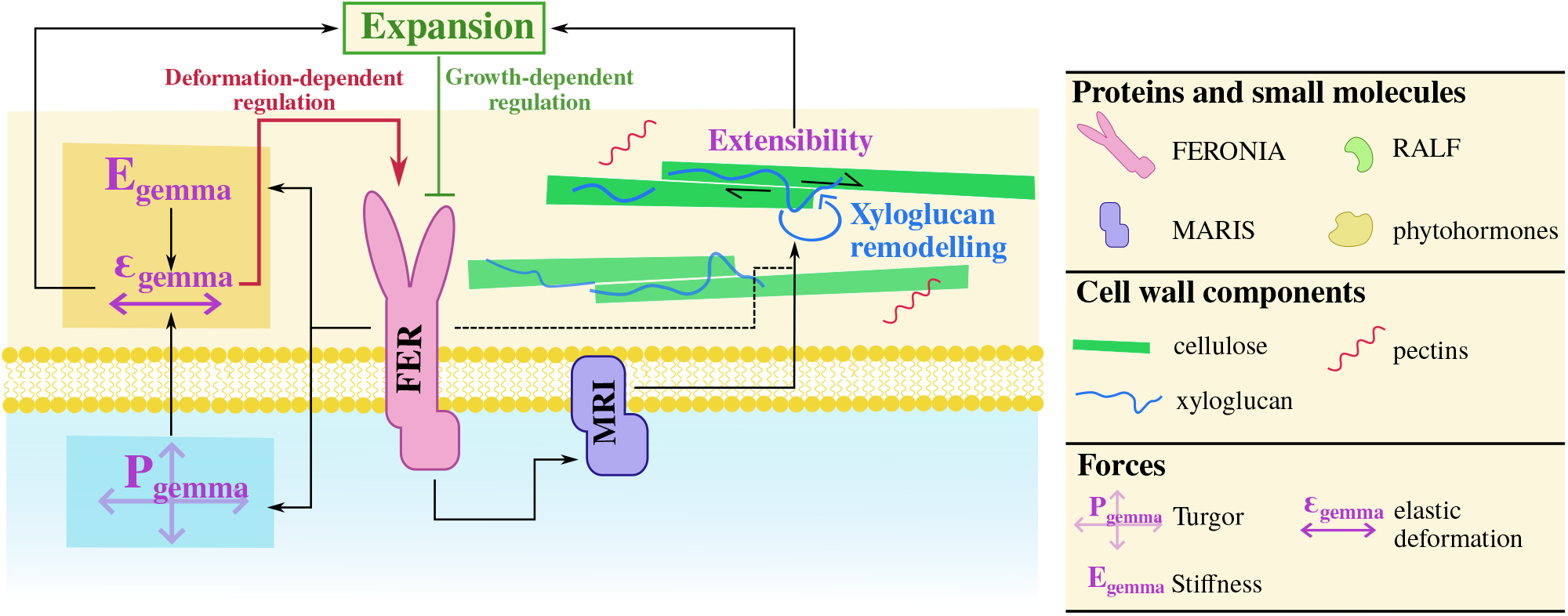
Molecular interpretation of the dual function of the FERONIA pathway. Dual role of FERONIA on mechanical parameters (left side of the schematics) and on the extensibility, notably through MARIS and xyloglucan remodelling (right side of the schematics). The twofold regulation of FERONIA by a positive deformation-dependent and a negative growth-dependent regulation is also represented.

Focusing on effectors directly affecting the cell wall, an transcriptomic analysis of *mri*-1 and *fer*-1 mutants (*42*) allowed us to hypothesise a role for XTH activity, which participates in xyloglucan remodelling, as a FER/MRI effector. Similarly, in At*fer* (*59*), down-regulated transcripts and proteins are enriched in the ‘cell wall related genes’, notably glycosyl hydrolase and in particular XTH. This hypothesis is supported by the alteration of xyloglucan contents of *fer*-2 gemmae, revealed by our immunolocalisation experiments. Reduced xyloglucan labelling in xyloglucan backbone biosynthesis mutants was associated with decrease in cell wall stiffness and in turgor but no growth phenotype (*60*), while xyloglucan side-chains biosynthesis mutants *xxt1/xxt2* associate xyloglucan reduction to growth and morphogenesis phenotypes, extensibility reduction (*45, 61*) but no significant effect on mechanical properties (*62*). Overexpression of Arabidopsis xyloglucan endotransglucosylase/hydrolase leads to a stimulation of hypocotyl growth and extensibility as well as effects on cell wall elasticity and deposition (*43*). Therefore, xyloglucan remodelling may account for changes in cell wall mechanics in *fer*-2 and is a plausible explanation of extensibility changes, potentially leading to dual role of *FERONIA* in mechanics and growth. The observed effect of *FERONIA* on xyloglucan content in the cell wall leads to the hypothesis that *FERONIA* might mediate compensatory mechanism between matrix polysaccharides. Indeed, it was shown that FERONIA activation can be mediated by pectin sensing (*18*), possibly enabling the compensation of pectin defects by xyloglucan.

We developed a mathematical model of the regulation of extensibility by the FER/MRI pathway, that recapitulates the main growth phenotypes of WT and *fer*-2. The model is based on the assumption of a double regulation of extensibility by a positive response to deformation and by a negative response to growth (Fig. 7). The assumption of a response to deformation is reinforced by the mechanical response of gemmae to a long term osmotic treatment. Indeed, the elastic modulus *E*_*gemma*_ and the osmotic pressure Π_*gemma*_ in *fer*-2 are lower than in WT under control growth condition but similar with +100 mM osmotic treatment, which corresponds to low deformation *ℰ*_*gemma*_. In Arabidopsis, FERONIA is inactivated under salt stress (*63*), suggesting an activation of FERONIA by deformation.

We now discuss the possible mechanistic basis of the double regulation of extensibility by FERONIA. We may first consider two distinct activation mechanisms upstream of FERONIA. Indeed, the response to cell wall deformation could originate in the activation of FERONIA by the cell wall status, by binding to pectins either directly through the malectin domain (*18, 19*) or indirectly through leucine-rich repeat extensins (*10*). We may then hypothesise that the response to cell wall expansion originate in RALF sensing (*21*). This would be possible if RALF synthesis depended on cell growth or if dilution of RALF by expansion were sensed by FERONIA, for instance. However, it is unclear whether pectin and RALF sensing can be uncoupled. It was hypothesised that RALF and pectin signals interact together through liquid-liquid phase separation based on formation of RALF-pectin condensate that act as stress sensor and promotes FERONIA clustering and massive endocytosis (*64*). Structural roles of some RALFs in the cell wall might also underlie a role of RALF as another mechanical sensor, and so RALF would rather mediate response to deformation (*65*). Therefore we may second consider that the sensitivities to elastic deformation and to growth are properties of the signalling pathway downstream of FERONIA, which would filter static (deformation) and dynamic (expansion) signals. Indeed, we can notice that rapid response of FERONIA is associated with growth arrest as apoplasmic pH increases or ROS increases (*5*), while longer-term regulation through transcriptional regulation may be sensitive to cell expansion. This is well illustrated by RALF1-induced root growth arrest in Arabidopsis, which involves rapid (~min) alkanisation of the apoplasm and slow (~h) auxin synthesis (*66*).

Individual-to-individual (or organ-to organ) variability differs in extent and in potential function according to context. Seed-to-seed variability in germination is relatively large, associated with a bet-hedging strategy to survive in an unpredictable environment (*67*), whereas sepal-to-sepal variability in morphology is relatively small, potentially associated with their function in flower protection (*53*). Here we found than the FER/MRI pathway is involved in limiting growth variability, like FERONIA in Arabidopsis roots (*15*). Indeed, the regulatory network underlying our model comprises a *FERONIA*-dependent negative feedback on growth; such negative autoregulation is known to promote robustness to external perturbations (*68*). We found that, in reference culture conditions, *FERONIA* restricts variability in germination time, which we ascribed to the spatiotemporal averaging of gemma growth, similar to Arabidopsis sepals for which spatiotemporal averaging yields robust final shape (*53*). In the shoot apical meristem of Arabidopsis, it was also found that reducing the response to mechanical force, by reducing microtubule alignment, amplifies the variability of cell growth rate, which increases spatial averaging and so shape robustness (*52*). Altogether, we have provided a quantitative framework that we expect to be broadly applicable to the analysis of mechanical responses during development.

## Author contributions

E.M.: conceptualization, methodology, software, validation, formal analysis, investigation, writing-original draft preparation, visualization. M.R.: conceptualization, methodology, validation, investigation, writing-original draft preparation, visualization, funding acquisition. G.D.: validation, investigation. V.L.: software. L.G.: conceptualization, formal analysis. A.L.: conceptualization, methodology, funding acquisition. S.D.:conceptualization, methodology, writing-original draft preparation, project administration, funding acquisition. A.B.: conceptualization, methodology, formal analysis, writing-original draft preparation, project administration, funding acquisition.

## Data and materials availability

Data and software will be made available in public repositories.

## Supplementary materials

Materials and Methods Supplementary Text Figs. S1 to S5

Tables S1 to S4

References (*66-76*)

## Supplementary Materials for

## Materials and Methods

### Plant growth and mechanics: experimental setup

#### Plant culture

Mother plants of the gemmae are grown at 22 °C and 4600-5100 lx continuous light in a growth cabinet (Aralab) on solid Gamborg 1/2 (Gamborg B5 medium, Duchefa with 1.2% agar, Duchefa), with additional 20 mM of mannitol (sigma-aldrich) for the *mri*-1 plants and the associated WT *Marchantia polymorpha* subsp. ruderalis accessions: Takaragaike-1, Tak-1, (WT) and Takaragaike-2, Tak-2, (WT-2), or additional 1% sucrose (sigma-aldrich) for the *fer*-2 and *fer*-3 plants and the associated WT. Details on the used lines are available in Table S4 and corresponding genes in Table S3. For the dividing nuclei staining and immunostaining assays, plant were first grown on plates in liquid Gamborg 1/2.

#### Culture and mechanical assays in a chip

Every growth and mechanical assays of this study were made by cultivating gemmae up to 24 h, 8 h or 1 h in a microfluidic chip. Chips and plants are prepared and grown as described in (*37*), except that the surfactant (Tween 20) was diluted at 1/2000 instead of 1/1000. For continuous osmotic treatments, we added 100 mM of D-mannitol (Sigma-Aldrich) to the Gamborg 1/2 liquid medium.

To measure mechanical parameters, gemmae were grown for 1 h or 8 h in a microfluidic chips and deformed by successive osmotic steps of +100 mM, +250 mM and +900 mM of mannitol (Fig.3-A). Each step last 20 min which includes 10 min medium change (at rate 25 *µ L* /*min*) and 10 min of slower medium flow (at rate 8.33 *µ L* /*min*) to reach mechanical equilibrium.

For gemmae area measurements, image acquisition was performed in bright field at 16x magnification using a Zeiss Axiozoom V16, as in (*37*). For growth assays images were recorded every 30 min and for mechanical assays images were recorded every 30 second and the acquisition started 2 min before the first osmotic step.

### Statistical analyses

Statistical significance of the comparison between two samples was estimated using a non-parametric Wilcoxon test (the p-values are displayed and set to 0 when *<* 10^−3^). To estimate the size effect of the difference between two samples the RC (relative change: difference between the two median of the samples over the median of the reference sample) and the ES (effect size: median difference compared to the reference sample variability) were computed. For each experiment *n* corresponds to the number of gemmae and *rep*. to the number of independent chip experiments or independent experimental replicates for experiments not conducted in chips.

Variability is estimated thanks to the relative average absolute deviation (AAD). The relative AAD for a sample of median 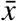 is given by equation S1.

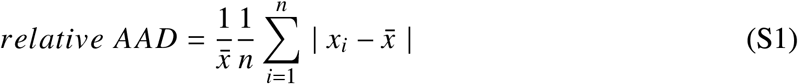

As samples present important size variation, the relative AAD for *T*_*start*_ and *G*_*eq*_ was computed on a random subset of values of size equal to 90% of the size of the smallest sample. The calculation was repeated on a number of random values subsets equal to the size of the subsets. For the local growth rate, a gemma AAD was computed on every individual over all local growth rate points. The median AAD over all individuals was then divided by median growth rate in order to avoid calculation error due to small local growth rate on individual gemmae.

Correlation analysis were performed using a Spearman’s correlation coefficient calculation and affine fits (for instantaneous growth rate *G*_*instant*_ correlation with *ℰ*_*gemma*_) are realised with the the lineregress function from the scipy.stats library.

### Image analysis

All image analysis were performed using python.

#### Area measurement

Image analysis to extract area evolution over time was performed as described in (*37*) although the size of the binary circular elements were adjusted for *fer*-2 as the image presented more irregularities (closing at 2.5 µm and opening at 20 µm).

#### Growth parameters computation

The two phases of growth are fitted by

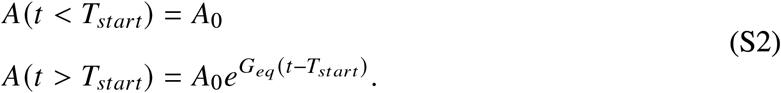

with *A* being gemma area. *T*_*start*_ and *G*_*eq*_ are obtained from successive fits as described in (*37*). For the correlation analysis, we used the instantaneous growth rate 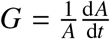, computed as in (*37*).

#### Mechanical parameters computation

We aimed at determining the average internal osmotic pressure of a gemmae Π_*gemma*_ as well as the bulk volumetric elastic modulus *E*_*gemma*_ defined by equation S3.

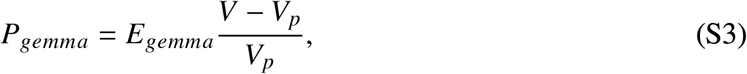

with *V* being the gemmae volume, *V*_*p*_ the plasmolysis volume and *P*_*gemma*_ the turgor pressure.

Gemmae are submitted to 4 osmotic concentrations: +0 mM, +100 mM, +250 mM and +900 mM. Plants remain turgid under the three first concentrations and are plasmolysed under 900 mM (Fig. 3-A). Mechanical parameters were deduced from the water potential Ψ equilibrium between the outside medium and the gemmae internal medium at each turgid pressure step.

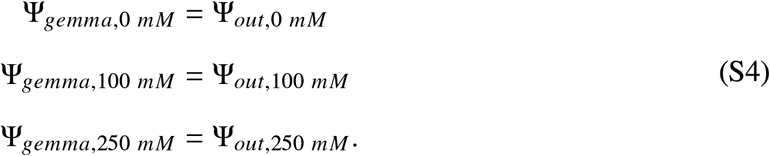

With the different component of the water potential: the hydrostatic pressure (the turgor pressure *T* for the internal medium and no over pressure for the outside media) and the osmotic pressure Π:

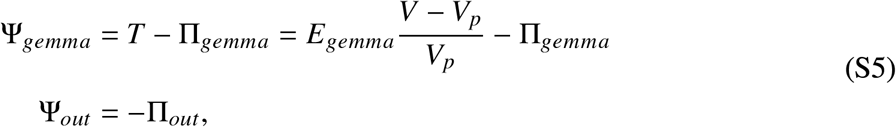

with *V*_*p*_ being the plasmolysis volume given by the +900 mM step. This yields

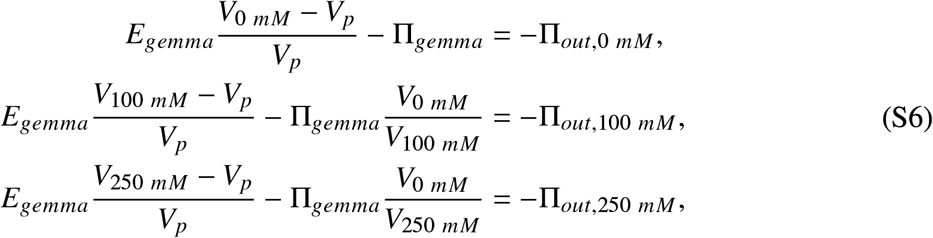

where 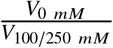 accounts for the dilution effects at each deformation step, as we hypothesized that osmoregulation does not occur on these time scales and therefore the number of osmolytes is constant.

Two equilibria are needed to deduce both the elastic modulus *E*_*gemma*_ and the turgor pressure Π_*gemma*_. As the gemmae exhibit a strain-stiffening behaviour, we estimate Π_*gemma*_ using the equilibrium closer to the plasmolysis (+100 mM and +250 mM) to stay in a similar strain regime.

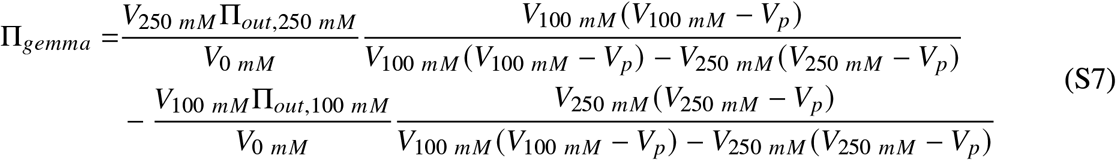

The turgor pressure *P*_*gemma*_ before the shocks is calculated according to

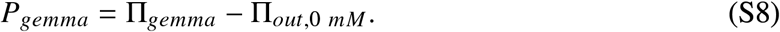

Finally, *E*_*gemma*_ is estimated at the equilibrium corresponding to the initial medium,

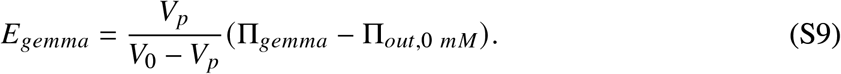

#### Measurement of local deformations

Local growth rate and local mechanical properties are assessed by first extracting local deformations on time courses of gemmae bright field images. Several processing steps are performed.

1. Only gemmae with two notches and the attachment point visible are selected. For the osmotic step strain rate calculation, a time point in each equilibrium state is selected manually.
2. The localisation of these three points (two notches and the attachment point) is detected manually for the first time point and is propagated to the next time points using contour fitting to adjust for movement and curvature to identify the notches (fig. S5-A step 1). *Details of the computation:* After binarisation of the image, the gemma contour is detected thanks to the openCV findContours function. High curvature points of this contour are calculated by fitting circles to contour portion and the highest one is extracted thanks to the find_peaks function of the scipy.signal library. Contours are aligned to the previous time point by minimizing the distance between the two contours and closest high sign peaks are associated.
3. Localisation of the points of interest are used to align gemmae according to the following procedure: the image is rotated to have the notch-to-notch axis horizontal and translated so that the attached point stays at the same coordinate over time. These operations are performed thanks to the openCV rotate and warpAffine functions (fig. S5-A step 1).
4. The optical flow, which is the apparent movement of the image objects, is computed on the processed image using the calcOpticalFlowFarneback function (openCV function applied to the image converted to gray scale with the following parameters pyr scale = 0.5, levels = 3, winsize = 15, iterations = 3, poly n = 5, poly sigma = 1.1, flags = cv.OPTFLOW_FARNEBACK_GAUSSIAN) which returns the *V*_*x*_ and *V*_*y*_ component of the velocity field (fig. S5-A step 2).
5. Background is removed using the same segmentation pathway as for area measurements.
6. To avoid noise caused by small movements on the image and to be able to detect tissue scale movement, the velocity field components are filtered using the ndimage.gaussian_filter of the scipy library with *σ* = 15*µm* (fig. S5-A step 3).
7. Local deformations are calculated as being the trace of the gradient i.e. 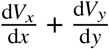 (fig. S5-A step 4).

To assess the reliability of the method, the global growth rate calculated based on area 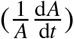 was compared to the sum of the local deformation or growth rate for each time step (fig. S5-B). Overall the local growth rate computed with optical flow is correlated with the global one, although it tends to be reduced. Moreover the dynamics over time look similar although noisier (fig. S5-C).

#### Local growth rate quantification

To compare local growth rate between gemmae, images were co-registered by aligning the notch-to-notch axis (thanks to the warpAffine function) and were scaled to the same area (using the openCV resize method). To visualise average patterns, median values of the local growth rate at a given localisation are plotted only for data point with a gemmae presence probability higher than 0.5. Local probability of gemmae presence was computed by assigning a probability of 1/*N*_*gemmae*_ to each aligned gemma and thresholding the obtained map to 0.5.

To quantify spatial patterning of the local growth rate, we proceeded first to a angular averaging. Each local growth rate field of a gemma (we work with resized field) is virtually dissected into 13 annuli centred at the mid distant between the two notches and with increased radius covering the whole gemmae once (see schematic in Fig. 2-E). The average local growth rate values was computed for each annulus.

For radial averaging local growth rate field are virtually dissected into 36 angular sectors (centred at the middle of the two notches, see schematic in Fig. 2-F). The average local growth rate value is computed for each sector.

#### Local mechanical parameters computation

Local displacement fields were also calculated at each equilibrium state of a given gemma. Gemmae were aligned and median local deformation was calculated. Due to boundary effect on mechanical parameters evaluation, mechanical parameters are evaluated only on the surface with a probability of gemmae presence higher than 0.7. A similar reasoning than for the global mechanical parameters is applied for local elastic modulus and pressures. If we do not account for the dilution effect and we note the deformation from the equilibrium i to 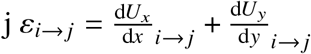, the local equilibrium equations are

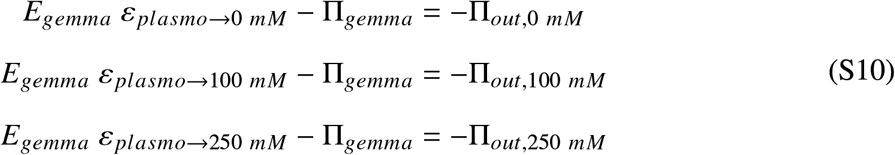

So we obtained the local Π_*gemma*_ as

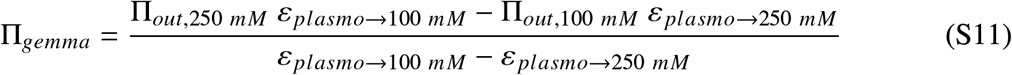

and so the local *E*_*gemma*_ is

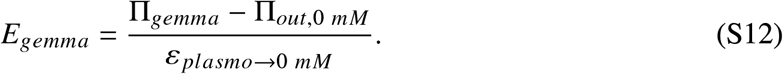

### Confocal microscopy

#### Nuclei staining

Gemmae were collected from the mother plants and grown on liquid Gamborg 1/2 (Duchefa), in cell filters, in the growth cabinet for 0 h, 4 h, 8 h or 19 h. Gemmae in the cell filters were then soaked on Gamborg 1/2 with 10 µm EdU solution (from the Click-iT EdU Imaging Kits Protocol, ThermoFischer) for 2 hours, 22 °C under continuous light. The cell fixation, permeabilisation and EdU detection are made as described in (*16*) (using the Click-iT kit with Alexa Fluor 647 dye). An additional PBS wash step (four times) was added both at the end of the permeabilisation and detection phase. After soaking in chloral hydrate for 30 min, gemmae were washed four times in PBS and mounted in Immersol W 2010 oil solution (Zeiss) for imaging.

Imaging was performed using an inverted confocal microscope TCS SP8 (Leica) with the HC PL APO 20x/0,75 IMM CORR CS2 objective. The white light laser (Leica) laser was set to 647 nm for excitation (100%) and the HyD1 detector detection windows to 657-680 nm.

The number of nuclei was automatically detected using the ImageJ 3D Nuclei Segmentation plugin with the “MaxEntropy” algorithm for the automatic thresholding. The obtained segmented image was then projected using ImageJ. Nuclei number was assessed with the Analyze Particles tools by selecting big and round enough particles (size set over 8 *µm*^2^ and circularity over 0.2). Analysis was automated thanks to the following macro.

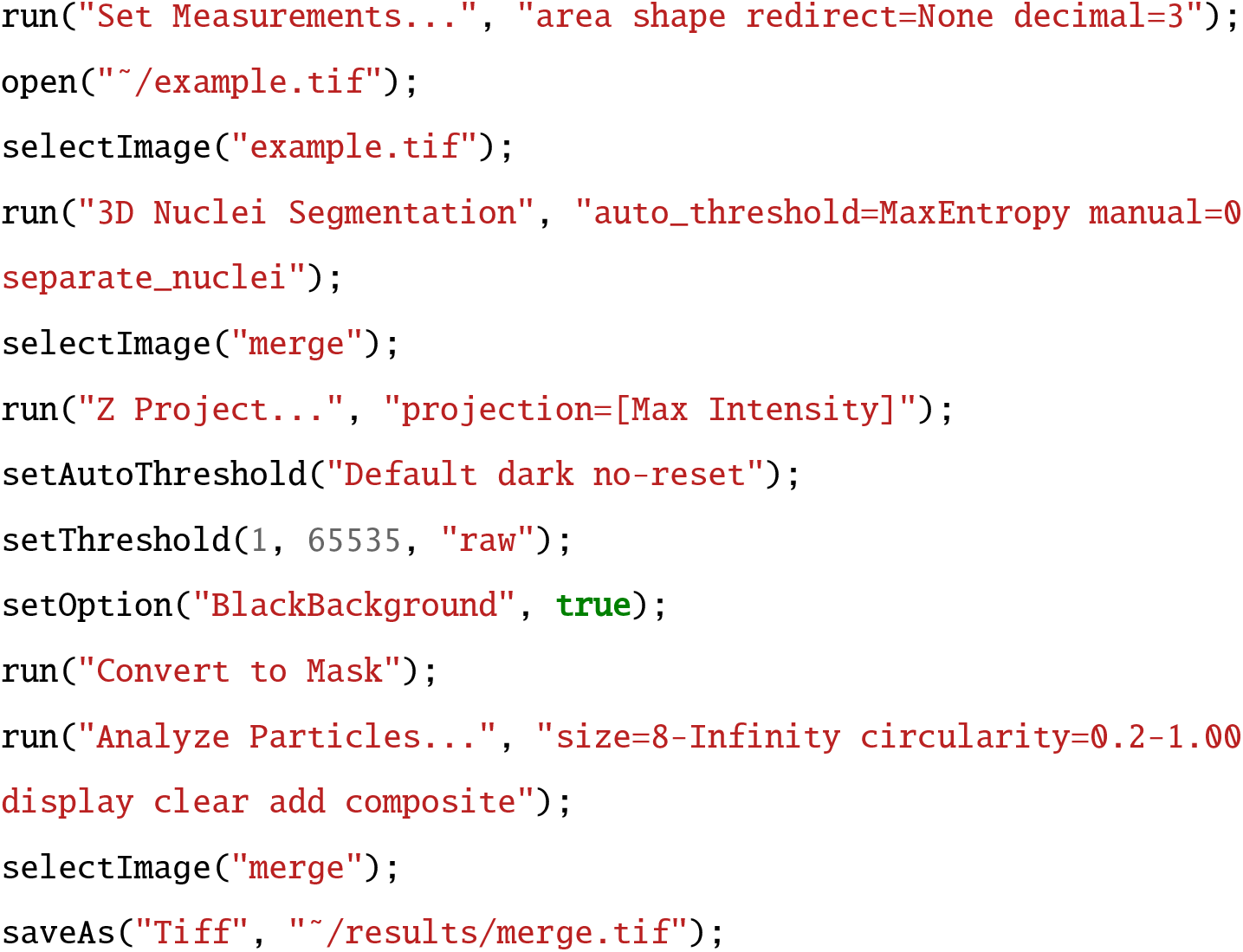

### Cell wall composition analysis

#### Resin Embedding and Immunofluorescence Labelling

Marchantia gemmae were collected and each condition (WT and *fer*-2) were processed with manual classical embedding adapted from (*69*) as follows. Samples were fixed for 1 h 30 in 1% paraformaldehyde and 1% glutaraldehyde mixture (v/v) in 0.1 M sodium cacodylate buffer (pH 7.2) and washed (5 min) in ultrapure water. Then, the samples were dehydrated in an ethanol series (30%, 50%, 70% and 100%) for 2 h each and embedded in LRW resin through a LRW—ethanol series (25%, 50%, 75% and 100%), for 24 h each. Finally, they were embedded (24 h) in LRW resin complemented with the UV catalyst benzoin methyl ether (0.5% w/v) and polymerized for 48 h at 4°C with UV light. Sections from resin blocks (2 µm - EM UC6 Leica microsystems) were collected on 10 well slides previously coated with poly-L-lysine (0.01% v/v). The resin sections were blocked in PBS-Tween 20 0.1% (w/v) supplemented with 3% of BSA (bovin serum albumin) and NGS 1/20 (normal goat serum, v/v) for 30 min. Sections were washed in PBS-T + 1% BSA (5 x 5 min) and incubated overnight at 4°C in a wet chamber with the primary antibody (table S2). After washing in PBS-T + 1% BSA (5 x 5 min), the sections were incubated for 2 h at 25°C in a wet chamber with a rat secondary antibody coupled to Alexa 647 (d:1/200, In Vitrogen). Finally, they were washed in PBS-T + 1% BSA (5 x 5 min) and ultrapure water (2 x 5 min). Fluorescence on sections was observed using a macroscope Axiozoom Zeiss with a fluorescence filter set 50 (BP 640/30 BP 690/50), an exposure time of 400 ms and a range of 50 µm, slides: 11, interval: µm. Negative controls were performed by the omission of the primary antibody (table S2).

### Image Analysis and Statistical Test

Fluorescence intensity measurements were performed on a region of interest (ROI) containing the central area of gemma and used to obtain a mean of fluorescence intensity value of the ROI. To automate the measurements, an ImageJ macro was using and adapted of (*70*). It manages the opening of an image, the projection of average intensity along the Z axis and the retrieval and saving of intensities along a line in a result table. Z-average projection is used to smooth out cutting effects, as each point on the line is itself an average of a user-defined orthogonal segment.

## Supplementary Text

### A simple estimate for the germination characteristic time *τ*

By neglecting the dilution effect, which is reasonable to get an estimate of early growth dynamics, we get the following dynamics for *W* (equation S13),

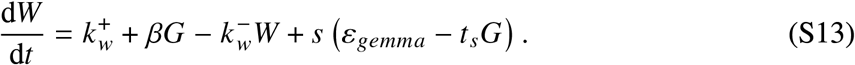

Thus *G* is given by:

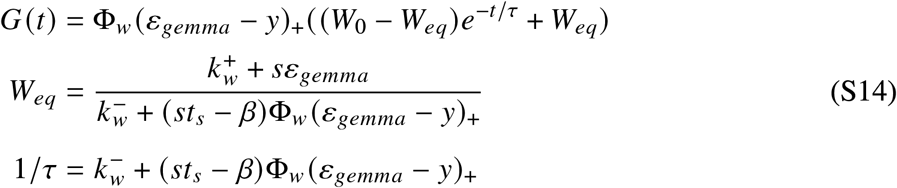

### Solution of the complete model for *W*

The complete model equation is given by

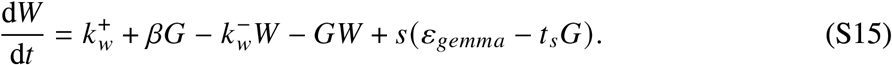

This can be solved analytically, as the right member of the equation can be re-organised as a second degree polynomial

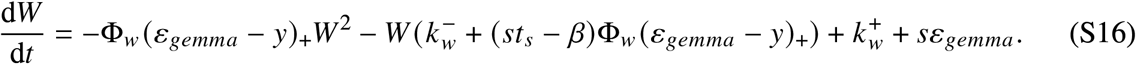

The roots of the polynomial, *w*_+_ and *w*_−_, are expressed are

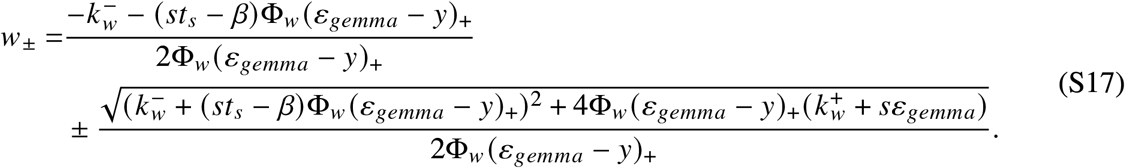

Equation S15 is thus equivalent to

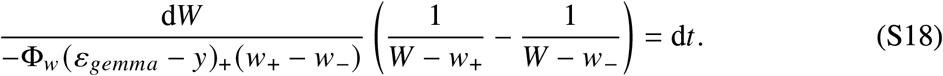

Integration of equation S18 gives the following expression for *W* (equation S19),

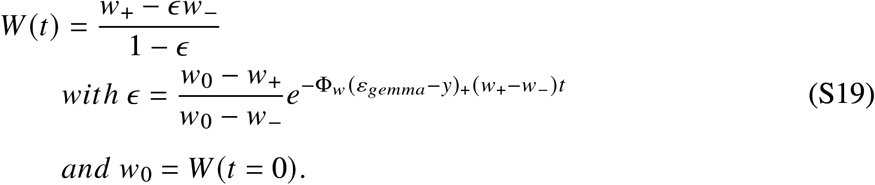

The equilibrium value for the elongation agent surface concentration is *W*_*eq*_ = *w*_+_. The equilibrium growth rate is thus given by

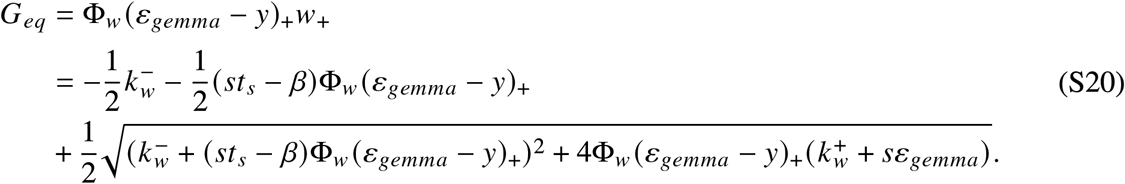

To characterise germination, we define a germination half-time *τ*_1/2_ so that 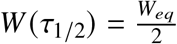, which gives

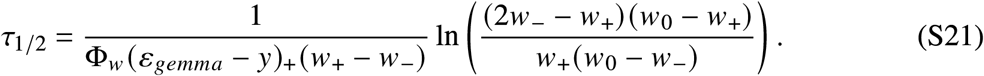

Estimates of the parameter values are given in Table S1.

### Variability estimates given by the model

We consider small fluctuations of the remodelling agent concentration *W* around an equilibrium value *W*_*eq*_, which is solution of Equation (S13), so that *W* = *W*_*eq*_ +Δ*W* and Δ*W* ≪ *W*_*eq*_, we linearise Equation S15 and get

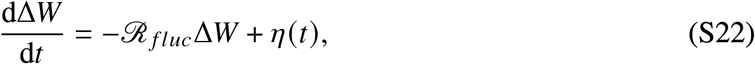

where

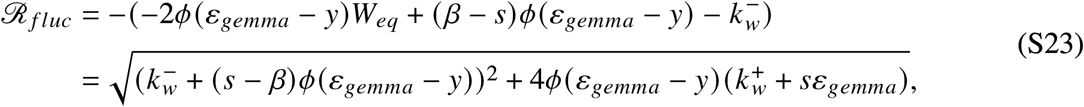

and *η* is a Gaussian white noise of amplitude *D*_*η*_.

We therefore predict

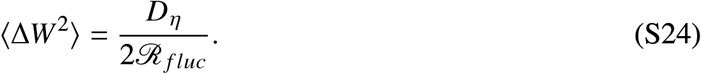

This gives the following relative fluctuation amplitude for *G*_*eq*_

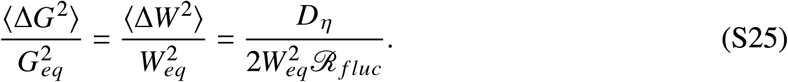

To evaluate the germination half-time fluctuation, we are interested in the sensitivity of the germination time to the initial value of *W* (*w*_0_), which is defined by

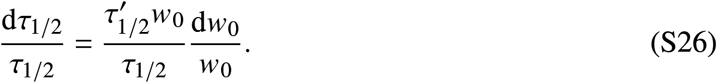

So the sensitivity of the relative germination time to the initial value of *W* is obtained from Equation S21 as

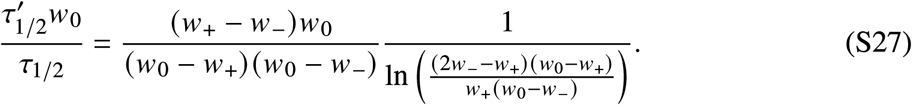

**Figure S1:**
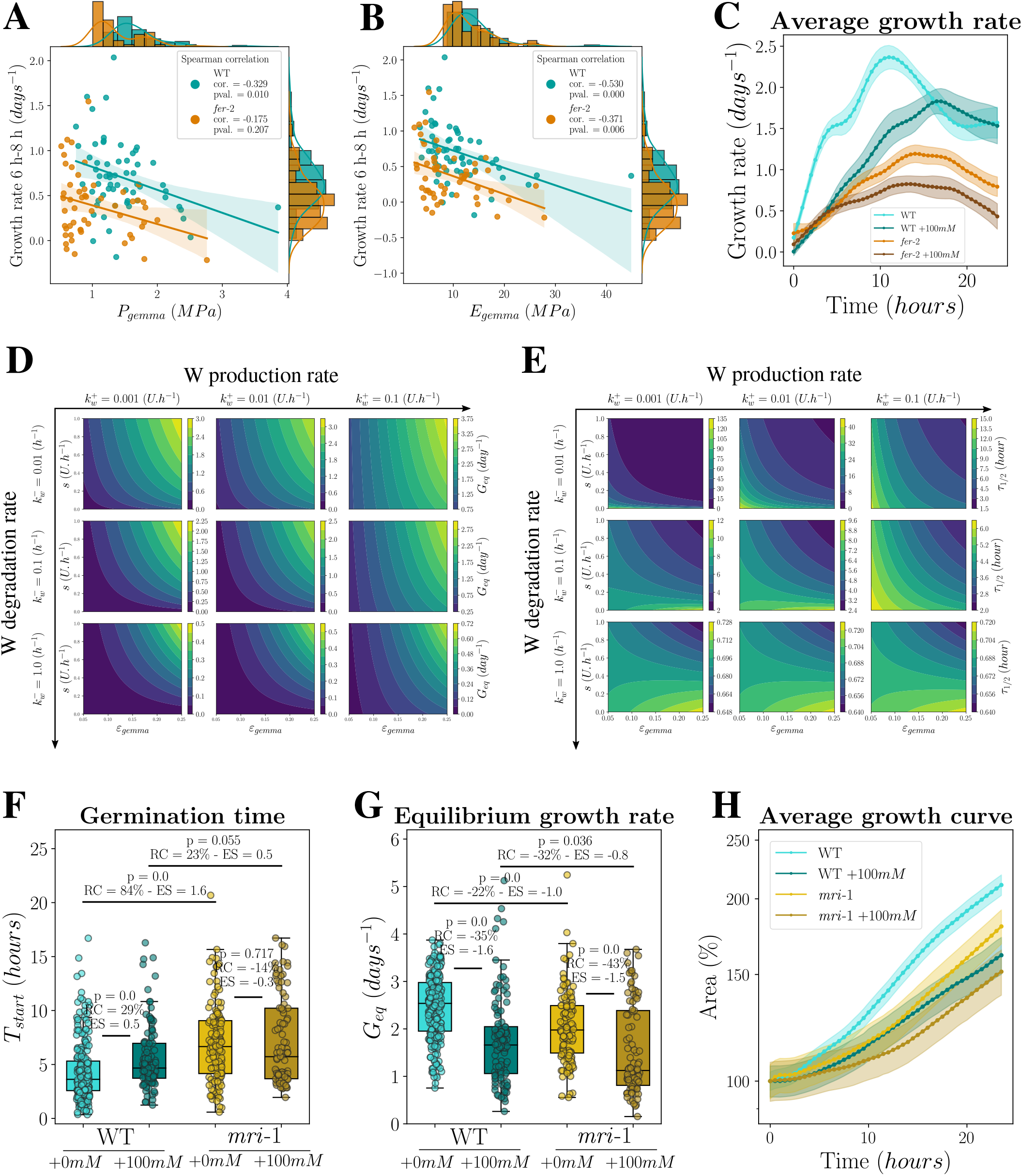
FERONIA and MARIS regulates extensibility. (**A-B**) Correlation analysis between instantaneous growth rate averaged between 6 h and 8 h post imbibition and mechanical parameters (**A**) turgor *P*_*gemma*_ and (**B**) elastic modulus *E*_*gemma*_ for both WT and *fer*-2. Scatter plot and regression line with 95% confident interval are represented as well as histograms per genotype for each variable. *cor*. stands for the Spearman correlation coefficient. WT: n = 62, rep. = 3 and *fer*-2: n = 56, rep. = 3. (**C**) Average growth rates for WT and *fer*-2, with and without osmotic treatment (+100 mM or +0 mM mannitol). Shaded areas correspond to the 95% confident interval. WT(+0 mM): n = 193, rep. = 4. WT(+100 mM): n = 136, rep. = 3. *fer*-2(+0 mM): n = 69, rep. = 6. *fer*-2(+100 mM): n = 70, rep. = 3. (**D-E**) Parameters space exploration for the mathematical model for different values of the W production rate 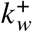 and degradation rate 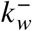. Model output according to the sensing parameter *s* and the elastic deformation *ℰ*_*gemma*_ for (**D**) the equilibrium growth rate *G*_*eq*_ and (**E**) the germination characteristic time *τ*_1/2_. Other parameters are set to: *t*_*s*_ = 1 *h, β* = 0.5 *U, E* = 10 *MPa* and *Y* = 0.01. (**F-H**) Parametrisation of growth of WT and WT-2 as well as of *mri*-1 for plants grown in reference medium with +0 mM or with +100 mM mannitol. (**F**) Box plot and scatter plot of the germination starting time *T*_*start*_. (**G**) Box plot and scatter plot of the equilibrium growth rate *G*_*eq*_. (**H**) Average area over time (relative to initial area). Shaded areas are the 95% confident interval. WT: n = 238, rep. = 7, WT(+100 mM): n = 109, rep. = 5, *mri*-1: n = 121, rep. = 4, *mri*-1(+100 mM): n = 93, rep. = 3.

**Figure S2:**
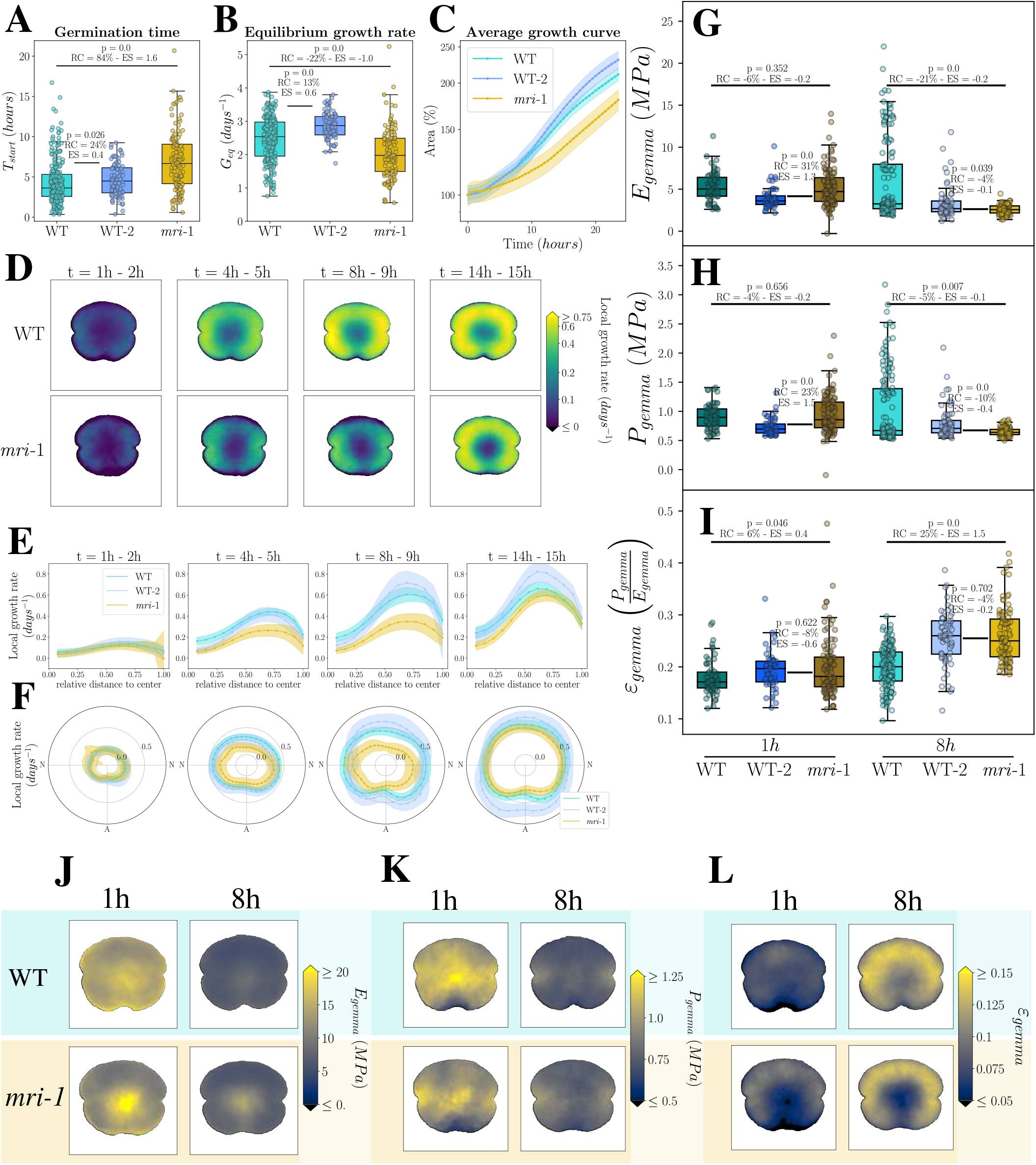
*mri*-1 mutant diplays a growth phenotype similar to *fer*-2 but does not have a mechanical parameters phenotype. (**A-C**) Parametrisation of growth of WT and WT-2 as well as of *mri*-1. (**A**) Box plot and scatter plot of the germination starting time *T*_*start*_. (**B**) Box plot and scatter plot of the equilibrium growth rate *G*_*eq*_. (**C**) Average area over time (relative to initial area). Error bars are the 95% confident interval. WT: n = 238, rep. = 7. WT-2: n = 84, rep. = 3. *mri*-1: n = 121, rep. = 3. (**D**) Median local growth rate maps for WT and *mri*-1. The median local growth rate is calculated over the individuals and over one hour at different time intervals (1 h-2 h, 4 h-5 h, 8 h-9 h and 14 h-15 h after imbibition). It is represented with a symmetric logarithmic color scale. WT: n = 96, rep. = 4. *mri*-1: n = 70, rep. = 4. (**E-F**) Radial and circumferential quantification of the local growth rate for WT, WT-2 and *mri*-1. (**E**) Mean local growth rate of the gemma surface at a given distance from the centre of the gemmae, averaged over all orientations. (**F**) Mean local growth rate of the gemma surface in a given angular sector, averaged over all distances to the centre. Shaded areas are the 95% confident interval. WT: n = 96, rep. = 4. WT-2: n = 50, rep. = 3. *mri*-1: n = 70, rep. = 4. (**G-I**) Box plot and scatter plot of measured mechanical properties at the scale of the whole gemma for WT, WT-2 and *mri*-1 gemmae at 1 h and 8 h after imbibition. (**G**) Volumetric elastic modulus *E*_*gemma*_, (**H**) turgor pressure *P*_*gemma*_ and (**I**) elastic deformation *ℰ*_*gemma*_ = *P*_*gemma*_/*E*_*gemma*_. WT: n(1 h) = 61, n(8 h) = 147, rep. = 3. WT-2: n(1 h) = 53, n(8 h) = 65, rep. = 2. *mri*-1: n(1 h) = 104, n(8 h) = 89, rep. = 3. (**J-L**) Local maps of local mechanical parameters for WT and *mri*-1 gemmae at 1 h and 8 h after imbibition, with linear color scale. Parameters are estimated locally from the median displacement during the osmotic steps. (**J**) Maps of the local elastic modulus *E*_*gemma*_, (**K**) maps of the turgor *P*_*gemma*_ and (**L**) maps of the elastic deformation, *ℰ*_*gemma*_. WT: n(1 h) = 54, n(8 h) = 81, rep. = 2. *mri*-1: n(1 h) = 52, n(8 h) = 82, rep. = 2.

**Figure S3:**
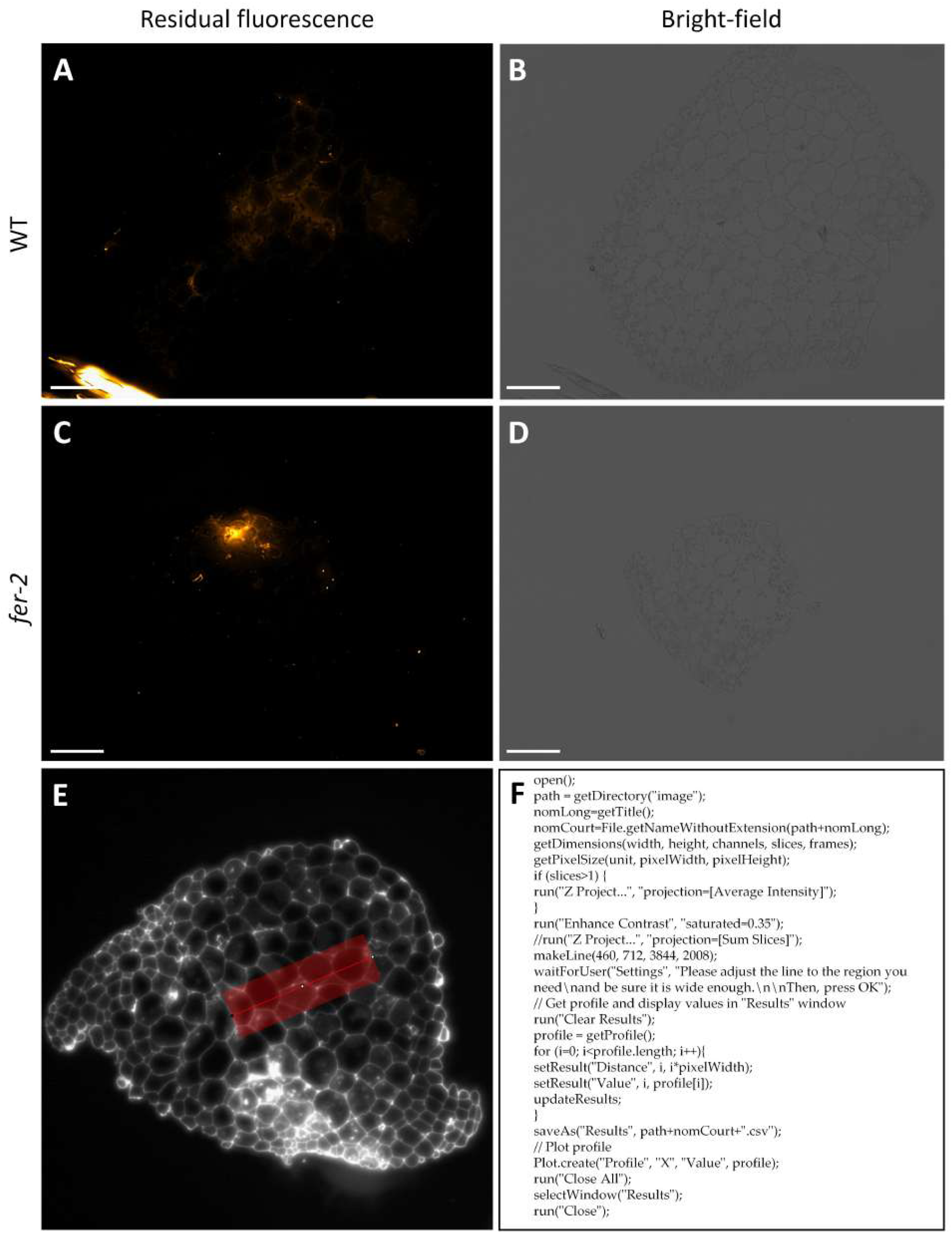
Immunolabelling negative controls of Marchantia gemmae after embedding and region of interest (ROI) defined for each pictures of Marchantia and ImageJ macro. Negative controls samples in which primary antibodies were omitted treated only with secondary antibody coupled Alexa647 show no fluorescence in (**A**,**B**) WT and (**C**,**D**) *fer*-2 gemmae. (**E**) The red area corresponds to the fluorescent measurement area. (**F**) To automate the fluorescent measurements, an ImageJ macro developed for fluorescence measurements. Scale bars are 100 µm.

**Figure S4:**
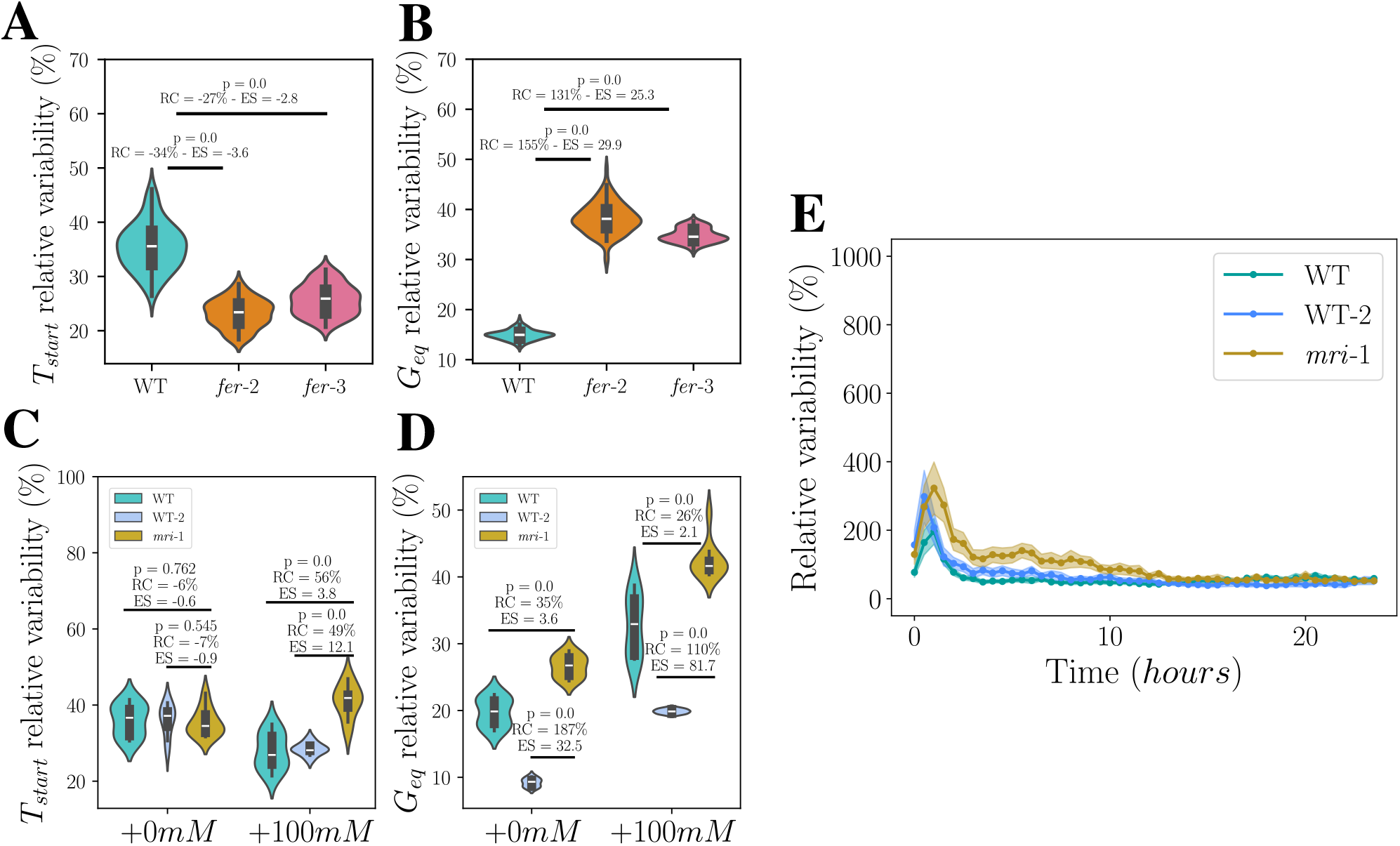
*FERONIA* and *MARIS* regulates growth variability. (**A-B**) Relative variability estimated by relative average absolute deviation of the experimental growth parameters for *fer*-2, *fer*-3 and WT for (**A**) the germination time *T*_*start*_ and (**B**) the equilibrium growth rate *G*_*eq*_. WT: n = 178 and rep. = 5. *fer*-2: n = 141 and rep. = 3. *fer*-3: n = 104 and rep. = 3. (**C-D**) Relative variability estimated by relative average absolute deviation of the experimental growth parameters for WT, WT-2 and *mri*-1 at +0 mM and +100 mM for (**C**) the germination time *T*_*start*_ and (**D**) the equilibrium growth rate *G*_*eq*_. WT: n = 238, rep. = 7. WT-2: n = 84, rep. = 3. *mri*-1: n = 121, rep. = 3. (**E**) Quantification of the variability (AAD over the median) of the local growth rate during 24h of growth, for WT, WT-2 and *mri*-1. Shaded areas represent the 95% confident interval. WT: n = 96, rep. = 4. WT-2: n = 50, rep. = 3. *mri*-1: n = 70, rep. = 4.

**Figure S5:**
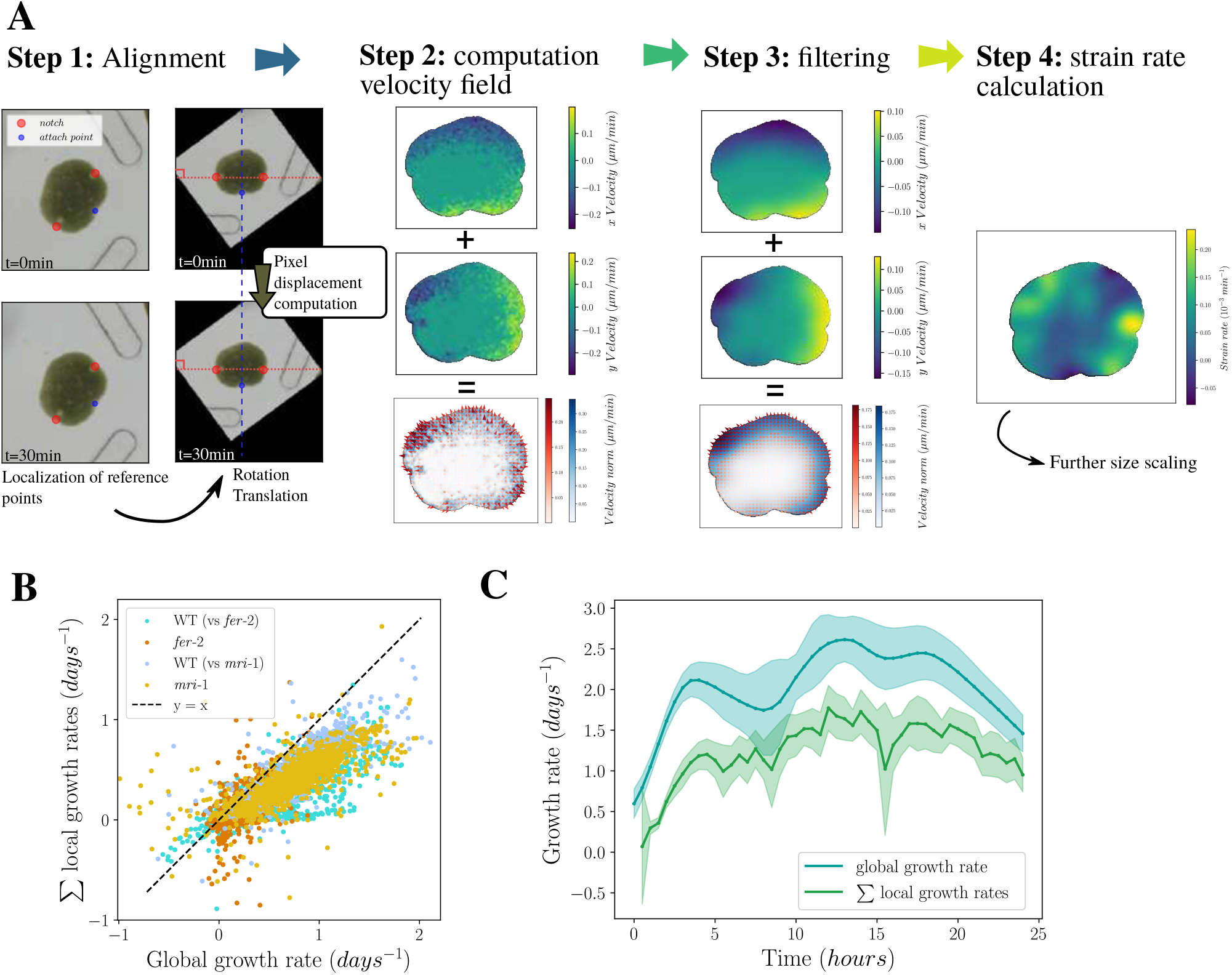
Method to compute to local displacements (to obtain local deformation or local growth maps). (**A**) Steps of the strain rate computation method, details in the method. (**B**) Comparison of the global growth rate measured by 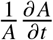 and the sum of the local growth rates during a time step *∂t* for growth experiments in chip with different genotypes (WT and *fer*-2 with mother plant on gamborg 1/2 supplemented with 1% sucrose, WT and *mri*-1 with mother on gamborg 1/2 supplemented with 20 mM mannitol). WT (vs *fer*-2): n = 61, rep. = 3. *fer*-2: n = 57, rep. = 3. WT (vs *mri*-1): n = 96, rep. = 4. *mri*-1: n = 70, rep. = 4. (**C**) Comparison of the global growth rate dynamics and the sum of the local growth rates for a given WT growth experiment in a chip. n = 21, rep. = 1. Shaded areas represent the 95% confident interval.

**Table S1:** Parameters estimates for the model.

| Parameter | Estimated value (unit) | References |
| --- | --- | --- |
| P - turgor pressure | 0.5-2.5 (MPa) | this work (osmotic steps measurements) |
| Y - yielding threshold | few | this work (deduction from growth arrest) |
| E - elastic modulus | 2-20 (MPa) | this work (osmotic steps measurements) |
| $\Phi_w$ - molecular extensibility | $10^{-5} (s^{-1} \cdot MPa^{-1}) \cdot U^{-1}$ | (71) |
| $k_w^+$ - W production rate | $(U \cdot h^{-1})$ | arbitrary units |
| $k_w^-$ - W degradation rate | $(hours^{-1} \text{ to } days^{-1})$ | (72) |

**Table S2:** Details of the epitopes recognised by the monoclonal antibodies used in the study.

| Cell Wall Glycomolecules |  | mAb | Epitope | References |
| --- | --- | --- | --- | --- |
| Hemicellulose | Xyloglucan | LM15 | Xylosylated xyloglucan (XXXG) | (73) |
|  |  | LM24 | Galactosylated xyloglucan (XXLG) | (74) |
|  |  | LM25 | Galactosylated xyloglucan (XXLG, XLLG) | (74) |
| | Heteromannan | LM21 | $\beta - (1 \rightarrow 4)$-mannan backbone epitope from DP2 to DP5 | (75) |

**Table S3:** Genes of interest.

| Gene name (abbreviation) | Gene ID |
| --- | --- |
| *FERONIA* (FER) | Mp4g15890, Mapoly0869s0001 |
| *MARIS* (MRI) | Mp7g17560, Mapoly0051s0094.1 |

**Table S4:** Marchantia lines.

| Genotype | Description | WT back-ground | Origin |
| --- | --- | --- | --- |
| Tak-1 | WT male Marchantia | - | - |
| Tak-2 | WT female Marchantia (WT-2) | - | - |
| *mri-1* | *MARIS* knock-down - T-DNA inserted in the promotor region | Tak-1 x Tak-2 | (33), (32) |
| *fer-2* | *FERONIA* knock-out Crispr/Cas mediated insertion | Tak-1 | (76) |
| *fer-3* | *FERONIA* knock-out Crispr/Cas mediated insertion | Tak-1 | (76) |

## References and Notes

1. D. J. Cosgrove, Catalysts of Plant Cell Wall Loosening. F1000Research 5, F1000 Faculty Rev–119 (2016), doi:10.12688/f1000research.7180.1.

2. O. Hamant, J. Traas, The Mechanics behind Plant Development. New Phytologist 185 (2), 369–385 (2010), doi:10.1111/j.1469-8137.2009.03100.x.

3. D. J. Cosgrove, Structure and Growth of Plant Cell Walls. Nature Reviews Molecular Cell Biology pp. 1–19 (2023), doi:10.1038/s41580-023-00691-y.

4. O. Hamant, E. S. Haswell, Life behind the Wall: Sensing Mechanical Cues in Plants. BMC Biology 15 (1), 59 (2017), doi:10.1186/s12915-017-0403-5.

5. X. Zhang, Z. Yang, D. Wu, F. Yu, RALF–FERONIA Signaling: Linking Plant Immune Response with Cell Growth. Plant Communications 1 (4) (2020), doi:10.1016/j.xplc.2020.100084.

6. S. Li, Y. Zhang, To Grow or Not to Grow: FERONIA Has Her Say. Molecular Plant 7 (8), 1261–1263 (2014), doi:10.1093/mp/ssu031.

7. J.-M. Escobar-Restrepo, et al., The FERONIA Receptor-like Kinase Mediates Male-Female Interactions During Pollen Tube Reception. Science 317 (5838), 656–660 (2007), doi:10.1126/science.1143562.

8. F. Yu, et al., FERONIA Receptor Kinase Controls Seed Size in Arabidopsis Thaliana. Molecular Plant 7 (5), 920–922 (2014), doi:10.1093/mp/ssu010.

9. M. Haruta, G. Sabat, K. Stecker, B. B. Minkoff, M. R. Sussman, A Peptide Hormone and Its Receptor Protein Kinase Regulate Plant Cell Expansion. Science 343 (6169), 408–411 (2014), doi:10.1126/science.1244454.

10. K. Dünser, et al., Extracellular Matrix Sensing by FERONIA and Leucine-Rich Repeat Extensins Controls Vacuolar Expansion during Cellular Elongation in Arabidopsis Thaliana. The EMBO Journal 38 (7), e100353 (2019), doi:10.15252/embj.2018100353.

11. H. Guo, et al., Three Related Receptor-like Kinases Are Required for Optimal Cell Elongation in Arabidopsis Thaliana. Proceedings of the National Academy of Sciences 106 (18), 7648–7653 (2009), doi:10.1073/pnas.0812346106.

12. S. D. Deslauriers, P. B. Larsen, FERONIA Is a Key Modulator of Brassinosteroid and Ethylene Responsiveness in Arabidopsis Hypocotyls. Molecular Plant 3 (3), 626–640 (2010), doi: 10.1093/mp/ssq015.

13. S. A. Kessler, et al., Conserved Molecular Components for Pollen Tube Reception and Fungal Invasion. Science 330 (6006), 968–971 (2010), doi:10.1126/science.1195211.

14. Q. Duan, D. Kita, C. Li, A. Y. Cheung, H.-M. Wu, FERONIA Receptor-like Kinase Regulates RHO GTPase Signaling of Root Hair Development. Proceedings of the National Academy of Sciences 107 (41), 17821–17826 (2010), doi:10.1073/pnas.1005366107.

15. H.-W. Shih, N. D. Miller, C. Dai, E. P. Spalding, G. B. Monshausen, The Receptor-like Kinase FERONIA Is Required for Mechanical Signal Transduction in Arabidopsis Seedlings. Current Biology 24 (16), 1887–1892 (2014), doi:10.1016/j.cub.2014.06.064.

16. M. A. Mecchia, et al., The Single Marchantia Polymorpha FERONIA Homolog Reveals an Ancestral Role in Regulating Cellular Expansion and Integrity (2022), doi:10.1101/2020.12.23.424085.

17. K. Hématy, et al., A Receptor-like Kinase Mediates the Response of Arabidopsis Cells to the Inhibition of Cellulose Synthesis. Current Biology 17 (11), 922–931 (2007), doi:10.1016/j.cub.2007.05.018.

18. W. Feng, et al., The FERONIA Receptor Kinase Maintains Cell-Wall Integrity during Salt Stress through Ca2+ Signaling. Current Biology 28 (5), 666–675.e5 (2018), doi:10.1016/j.cub.2018.01.023.

19. W. Lin, et al., Arabidopsis Pavement Cell Morphogenesis Requires FERONIA Binding to Pectin for Activation of ROP GTPase Signaling. Current Biology 32 (3), 497–507.e4 (2022), doi:10.1016/j.cub.2021.11.030.

20. A. Y. Cheung, FERONIA: A Receptor Kinase at the Core of a Global Signaling Network. Annual Review of Plant Biology 75 (Volume 75, 2024), 345–375 (2024), doi: 10.1146/annurev-arplant-102820-103424.

21. D. Ji, T. Chen, Z. Zhang, B. Li, S. Tian, Versatile Roles of the Receptor-Like Kinase Feronia in Plant Growth, Development and Host-Pathogen Interaction. International Journal of Molecular Sciences 21 (21), 7881 (2020), doi:10.3390/ijms21217881.

22. A. Malivert, O. Hamant, Why Is FERONIA Pleiotropic? Nature Plants 9 (7), 1018–1025 (2023), doi:10.1038/s41477-023-01434-9.

23. N. F. Keinath, et al., PAMP (Pathogen-associated Molecular Pattern)-Induced Changes in Plasma Membrane Compartmentalization Reveal Novel Components of Plant Immunity *. Journal of Biological Chemistry 285 (50), 39140–39149 (2010), doi:10.1074/jbc.M110.160531.

24. L. Vaahtera, J. Schulz, T. Hamann, Cell Wall Integrity Maintenance during Plant Development and Interaction with the Environment. Nature Plants 5 (9), 924–932 (2019), doi:10.1038/s41477-019-0502-0.

25. A. Malivert, et al., FERONIA and Microtubules Independently Contribute to Mechanical Integrity in the Arabidopsis Shoot. PLOS Biology 19 (11), e3001454 (2021), doi:10.1371/journal.pbio.3001454.

26. N. Hervieux, et al., A Mechanical Feedback Restricts Sepal Growth and Shape in Arabidopsis. Current Biology 26 (8), 1019–1028 (2016), doi:10.1016/j.cub.2016.03.004.

27. O. Hamant, et al., Developmental Patterning by Mechanical Signals in Arabidopsis. Science 322 (5908), 1650–1655 (2008), doi:10.1126/science.1165594.

28. B. Bozorg, P. Krupinski, H. Jönsson, Stress and Strain Provide Positional and Directional Cues in Development. PLOS Computational Biology 10 (1), e1003410 (2014), doi:10.1371/journal.pcbi.1003410.

29. A. Creff, et al., Evidence That Endosperm Turgor Pressure Both Promotes and Restricts Seed Growth and Size. Nature Communications 14 (1), 67 (2023), doi:10.1038/s41467-022-35542-5.

30. H. Höfte, The Yin and Yang of Cell Wall Integrity Control: Brassinosteroid and FERONIA Signaling. Plant and Cell Physiology 56 (2), 224–231 (2015), doi:10.1093/pcp/pcu182.

31. A. Boisson-Dernier, C. M. Franck, D. S. Lituiev, U. Grossniklaus, Receptor-like Cytoplasmic Kinase MARIS Functions Downstream of CrRLK1L-dependent Signaling during Tip Growth. Proceedings of the National Academy of Sciences 112 (39), 12211–12216 (2015), doi:10.1073/pnas.1512375112.

32. J. Westermann, et al., An Evolutionarily Conserved Receptor-like Kinases Signaling Module Controls Cell Wall Integrity During Tip Growth. Current Biology 29 (22), 3899–3908.e3 (2019), doi:10.1016/j.cub.2019.09.069.

33. S. Honkanen, et al., The Mechanism Forming the Cell Surface of Tip-Growing Rooting Cells Is Conserved among Land Plants. Current Biology 26 (23), 3238–3244 (2016), doi:10.1016/j.cub.2016.09.062.

34. C. Liu, H. Yu, A. Voxeur, X. Rao, R. A. Dixon, FERONIA and Wall-Associated Kinases Coordinate Defense Induced by Lignin Modification in Plant Cell Walls. Science Advances 9 (10), eadf7714 (2023), doi:10.1126/sciadv.adf7714.

35. L. Bacete, et al., THESEUS1 Modulates Cell Wall Stiffness and Abscisic Acid Production in Arabidopsis Thaliana. Proceedings of the National Academy of Sciences 119 (1), e2119258119 (2022), doi:10.1073/pnas.2119258119.

36. H. Kato, Y. Yasui, K. Ishizaki, Gemma Cup and Gemma Development in Marchantia Polymorpha. New Phytologist 228 (2), 459–465 (2020), doi:10.1111/nph.16655.

37. V. Laplaud, E. Muller, N. Demidova, S. Drevensek, A. Boudaoud, Assessing the Hydromechanical Control of Plant Growth. Journal of The Royal Society Interface 21 (214), 20240008 (2024), doi:10.1098/rsif.2024.0008.

38. M. Shimamura, Marchantia Polymorpha : Taxonomy, Phylogeny and Morphology of a Model System. Plant and Cell Physiology 57 (2), 230–256 (2016), doi:10.1093/pcp/pcv192.

39. Materials and methods are available as supplementary material.

40. Y. Zhang, et al., Molecular Insights into the Complex Mechanics of Plant Epidermal Cell Walls. Science 372 (6543), 706–711 (2021), doi:10.1126/science.abf2824.

41. J. A. Lockhart, An Analysis of Irreversible Plant Cell Elongation. Journal of Theoretical Biology 8 (2), 264–275 (1965), doi:10.1016/0022-5193(65)90077-9.

42. T. O. Jobe, et al., An Omics Approach on Marchantia Polymorpha Single FERONIA and MARIS Homologs Confirms Links between Cell Wall Integrity and Abscisic Acid (2024), doi:10.1101/2024.11.26.625412.

43. E. Miedes, et al., Xyloglucan Endotransglucosylase/Hydrolase (XTH) Overexpression Affects Growth and Cell Wall Mechanics in Etiolated Arabidopsis Hypocotyls. Journal of Experimental Botany 64 (8), 2481–2497 (2013), doi:10.1093/jxb/ert107.

44. K. Ishida, R. Yokoyama, Reconsidering the Function of the Xyloglucan Endotransglucosylase/Hydrolase Family. Journal of Plant Research 135 (2), 145–156 (2022), doi:10.1007/s10265-021-01361-w.

45. Y. B. Park, D. J. Cosgrove, Changes in Cell Wall Biomechanical Properties in the Xyloglucan-Deficient Xxt1/Xxt2 Mutant of Arabidopsis. Plant Physiology 158 (1), 465–475 (2012), doi: 10.1104/pp.111.189779.

46. M. J. Peña, A. G. Darvill, S. Eberhard, W. S. York, M. A. O’Neill, Moss and Liverwort Xyloglucans Contain Galacturonic Acid and Are Structurally Distinct from the Xyloglucans Synthesized by Hornworts and Vascular Plants*. Glycobiology 18 (11), 891–904 (2008), doi: 10.1093/glycob/cwn078.

47. E. R. Rojas, S. Hotton, J. Dumais, Chemically Mediated Mechanical Expansion of the Pollen Tube Cell Wall. Biophysical Journal 101 (8), 1844–1853 (2011), doi:10.1016/j.bpj.2011.08.016.

48. R. Rollin, J.-F. Joanny, P. Sens, Physical Basis of the Cell Size Scaling Laws. eLife 12, e82490 (2023), doi:10.7554/eLife.82490.

49. A. Fruleux, S. Verger, A. Boudaoud, Feeling Stressed or Strained? A Biophysical Model for Cell Wall Mechanosensing in Plants. Frontiers in Plant Science 10 (2019).

50. N. Hervieux, et al., Mechanical Shielding of Rapidly Growing Cells Buffers Growth Heterogeneity and Contributes to Organ Shape Reproducibility. Current Biology 27 (22), 3468– 3479.e4 (2017), doi:10.1016/j.cub.2017.10.033.

51. A. Sampathkumar, et al., Subcellular and Supracellular Mechanical Stress Prescribes Cytoskeleton Behavior in Arabidopsis Cotyledon Pavement Cells. eLife 3, e01967 (2014), doi: 10.7554/eLife.01967.

52. M. Uyttewaal, et al., Mechanical Stress Acts via Katanin to Amplify Differences in Growth Rate between Adjacent Cells in Arabidopsis. Cell 149 (2), 439–451 (2012), doi:10.1016/j.cell.2012.02.048.

53. L. Hong, et al., Variable Cell Growth Yields Reproducible Organ Development through Spatiotemporal Averaging. Developmental Cell 38 (1), 15–32 (2016), doi:10.1016/j.devcel.2016.06.016.

54. X. Wang, et al., FERONIA Controls ABA-mediated Seed Germination via the Regulation of CARK1 Kinase Activity. Cell Reports 43 (11), 114843 (2024), doi:10.1016/j.celrep.2024.114843.

55. E. Li, G. Wang, Y.-L. Zhang, Z. Kong, S. Li, FERONIA Mediates Root Nutating Growth. The Plant Journal 104 (4), 1105–1116 (2020), doi:10.1111/tpj.14984.

56. D. Kierzkowski, et al., Elastic Domains Regulate Growth and Organogenesis in the Plant Shoot Apical Meristem. Science 335 (6072), 1096–1099 (2012), doi:10.1126/science.1213100.

57. H. Vogler, G. Santos-Fernandez, M. A. Mecchia, U. Grossniklaus, To Preserve or to Destroy, That Is the Question: The Role of the Cell Wall Integrity Pathway in Pollen Tube Growth. Current Opinion in Plant Biology 52, 131–139 (2019), doi:10.1016/j.pbi.2019.09.002.

58. C. M. Franck, et al., The Protein Phosphatases ATUNIS1 and ATUNIS2 Regulate Cell Wall Integrity in Tip-Growing Cells. The Plant Cell 30 (8), 1906–1923 (2018), doi:10.1105/tpc.18.00284.

59. P. Wang, et al., Integrated Omics Reveal Novel Functions and Underlying Mechanisms of the Receptor Kinase FERONIA in Arabidopsis Thaliana. The Plant Cell 34 (7), 2594–2614 (2022), doi:10.1093/plcell/koac111.

60. F. B. Daher, et al., Xyloglucan Deficiency Leads to a Reduction in Turgor Pressure and Changes in Cell Wall Properties, Affecting Early Seedling Establishment. Current Biology 34 (10), 2094–2106.e6 (2024), doi:10.1016/j.cub.2024.04.016.

61. E. E. Sowinski, et al., Lack of xyloglucan in the cell walls of the Arabidopsis xxt1/xxt2 mutant results in specific increases in homogalacturonan and glucomannan. The Plant Journal 110 (1), 212–227 (2022), doi:10.1111/tpj.15666.

62. F. Zhao, et al., Xyloglucans and Microtubules Synergistically Maintain Meristem Geometry and Phyllotaxis1 [OPEN]. Plant Physiology 181 (3), 1191–1206 (2019), doi:10.1104/pp.19.00608.

63. X. Liu, et al., FERONIA Coordinates Plant Growth and Salt Tolerance via the Phosphorylation of phyB. Nature Plants 9 (4), 645–660 (2023), doi:10.1038/s41477-023-01390-4.

64. M.-C. J. Liu, et al., Extracellular Pectin-RALF Phase Separation Mediates FERONIA Global Signaling Function. Cell 187 (2), 312–330.e22 (2024), doi:10.1016/j.cell.2023.11.038.

65. S. Schoenaers, et al., Rapid Alkalinization Factor 22 Has a Structural and Signalling Role in Root Hair Cell Wall Assembly. Nature Plants 10 (3), 494–511 (2024), doi: 10.1038/s41477-024-01637-8.

66. L. Li, et al., RALF1 Peptide Triggers Biphasic Root Growth Inhibition Upstream of Auxin Biosynthesis. Proceedings of the National Academy of Sciences 119 (31), e2121058119 (2022), doi:10.1073/pnas.2121058119.

67. K. Abley, R. Goswami, J. C. W. Locke, Bet-Hedging and Variability in Plant Development: Seed Germination and Beyond. Philosophical Transactions of the Royal Society B: Biological Sciences 379 (1900), 20230048 (2024), doi:10.1098/rstb.2023.0048.

68. U. Alon, An introduction to systems biology: design principles of biological circuits (Chapman and Hall/CRC) (2019).

69. I. Boulogne, et al., Biological and Chemical Characterization of Musa Paradisiaca Leachate. Biology 12 (10), 1326 (2023), doi:10.3390/biology12101326.

70. C. Mirande-Ney, et al., LAM2: An Unusual Laminaran Structure for a Novel Plant Elicitor Candidate. Biomolecules 13 (10), 1483 (2023), doi:10.3390/biom13101483.

71. H. Nonami, J. S. Boyer, Wall Extensibility and Cell Hydraulic Conductivity Decrease in Enlarging Stem Tissues at Low Water Potentials. Plant Physiology 93 (4), 1610–1619 (1990), doi:10.1104/pp.93.4.1610.

72. L. Li, et al., Protein Degradation Rate in Arabidopsis Thaliana Leaf Growth and Development. The Plant Cell 29 (2), 207–228 (2017), doi:10.1105/tpc.16.00768.

73. S. E. Marcus, et al., Pectic Homogalacturonan Masks Abundant Sets of Xyloglucan Epitopes in Plant Cell Walls. BMC Plant Biology 8 (1), 60 (2008), doi:10.1186/1471-2229-8-60.

74. H. L. Pedersen, et al., Versatile High Resolution Oligosaccharide Microarrays for Plant Glycobiology and Cell Wall Research *. Journal of Biological Chemistry 287 (47), 39429–39438 (2012), doi:10.1074/jbc.M112.396598.

75. J. J. Ordaz-Ortiz, S. E. Marcus, J. P. Knox, Cell Wall Microstructure Analysis Implicates Hemicellulose Polysaccharides in Cell Adhesion in Tomato Fruit Pericarp Parenchyma. Molecular Plant 2 (5), 910–921 (2009), doi:10.1093/mp/ssp049.

76. M. A. Mecchia, et al., Characterization of the Single FERONIA Homolog in Marchantia Polymorpha Reveals an Ancestral Role of CrRLK1L Receptor Kinases in Regulating Cell Expansion and Morphological Integrity. bioRxiv p. 2020.12.23.424085 (2020), doi:10.1101/2020.12.23.424085.

